# A conserved isoleucine gates the diffusion of small ligands to the active site of NiFe CO-dehydrogenase

**DOI:** 10.64898/2026.03.19.713016

**Authors:** Laura Opdam, Marta Meneghello, Chloé Guendon, Jade Chargelègue, Andrea Fasano, Aurore Jacq-Bailly, Christophe Léger, Vincent Fourmond

## Abstract

CO dehydrogenases (CODH) are metalloenzymes that reversibly oxidize CO to CO_2_, at a buried NiFe_4_S_4_ active site. The substrates, CO and CO_2_, need therefore to be transported through the protein matrix to reach the active site. The most likely pathway for intra-protein diffusion is the hydrophobic channel identified in the crystal structures. We used site-directed mutagenesis in an extensive manner to study the role of the highly conserved isoleucine 563 of *Thermococcus sp. AM4* CODH2. Certain substitutions significantly change the biochemical properties of the enzyme (K_M_ for CO, catalytic efficiency, product inhibition constant, catalytic bias…), and increase its resistance to the inhibitor O_2_, showing that isoleucine 563 plays a key role in determining access to the active site. The mutations have the same effects on the rates of binding of CO and O_2_ showing that the two molecules follow the same pathway and are not discriminated by the protein matrix. The I563F mutation decreases the bimolecular rate constant of inhibition by O_2_ 15-fold, and increases the IC50 20-fold, which is the strongest improvement in O_2_ resistance reported so far.

## Introduction

CO dehydrogenases (CODHs) are metalloenzymes that catalyze the oxidation of CO to CO_2_. They can be separated into two phylogenetically unrelated classes. The MoCu CODHs are members of the xanthine oxidase family and are found in aerobic bacteria, in which they support the aerobic growth on CO. They only work in the CO to CO_2_ direction^[1]^. On the contrary, the NiFe CODHs, which are the subject of this work, are found mostly in anaerobic organisms, where they catalyze the bidirectional conversion between CO and CO_2_ at a unique NiFe_4_S_4_ active site called the “C cluster”^[2,3]^. NiFe CODHs are known for their very high turnover frequencies (with rates approaching 90 000 s^-1^ reported for *Carboxydothermus hydrogenoformans* CODH2 at 105 °C^[4]^) and also very high catalytic efficiencies, as some of them, including CODH4 from *Carboxydothermus hydrogenoformans*, function at the diffusion limit over a large range of temperature^[5]^. The phylogenetic analysis of the available sequences of putative CODHs have further divided them into 8 clades^[6]^, although some of them are probably not CODHs but rather members of a newly discovered family of proteins of unknown function, whose active site contains iron, sulfur and oxygen, closely resembling that of the hybrid cluster proteins^[7,8]^.

CODHs are involved in a variety of cellular functions. Some form a tight complex with an acetyl-CoA synthase (ACS) subunit: CO is produced by the reduction of CO_2_ at the C cluster, and diffuses via an internal channel to the active site of the ACS subunit, in which it is added to a methyl group to form acetyl-CoA^[9,10]^. Bifunctional ACS/CODHs are involved in the Wood-Ljungdahl pathway of carbon fixation^[11]^. Other CODHs are coupled to hydrogenases to provide energy from the oxidation of CO to H_2_^[11]^ . Other proposed roles include the generation of reducing equivalents (NADPH)^[5,13]^ and the defence against oxidative stress^[5,13]^.

The C-cluster is a NiFe_4_S_4_ cluster organized as a NiFe_3_S_4_ distorted cubane in which one of the S ligands coordinates also an external Fe ion, called the “unique” or “exo” Fe^[14]^. The C-cluster is buried within the protein, so that the CO and CO_2_ molecules must diffuse through the protein matrix to reach the active site from the solvent.

In bifunctional ACS/CODH, the existence of an intramolecular channel connecting the C-cluster to the A-cluster (the site of acetyl-CoA synthesis in the ACS subunit) had been suspected from the observation that hemoglobin, a trap for CO, does not inhibit the activity of ACS/CODH^[15]^, and that only ^14^C-labeled acetyl-CoA is formed from ^14^CO_2_, even in the presence of a large excess of unlabeled CO^[9]^. The first crystallographic structure of *Moorella thermoacetica* ACS/CODH showed the presence of a 70 Å-long channel^[10]^. This channel was found to bind Xenon atoms, which is often considered evidence that a cavity is a functional gas channel^[16]^. Site-directed mutagenesis targeting some of the residues lining the channel had a strong effect on the enzyme reactivity^[17,18]^, showing that the channel is functional. However, while the path for intramolecular diffusion of CO had been identified, the pathway used by CO_2_ to reach the CODH active site was less clear. Using molecular dynamics, Wang and coworkers showed that CO_2_ diffuses within the enzyme by transiently occupying a series of cavities that are not visible in the crystal structure but rather formed dynamically^[19]^. They suggested that the reduction of CO_2_ to CO changes the hydrogen bonding network around the active site, rigidifying its surroundings and providing the directionality of the access to the C-cluster^[19]^. The recent structure of the ACS/CODH from *Clostridium autoethanogenum* has shown that the organization of the static channels is not conserved among the various ACS/CODHs^[20]^.

In the case of the other CODHs (which do not bind an ACS subunit), the nature of the tunnels has been less investigated so far. Hydrophobic channels could be observed in crystal structures; the first indication that these channels are functional came from the determination of the structure of *Ch* CODH2 inhibited by *n*-butyl-isocyanide, which occupies parts of the hydrophobic cavity^[21]^. More recently, Biester and coworkers exposed crystals of *Nitratidesulfovibrio vulgaris* Hildenborough (formerly *Desulfovibrio vulgaris* Hildenborough) CODH (hereafter *Nv* CODH) to high pressures of Xenon and showed that the channel in *Nv* CODH has two branches, one of which matches the beginning of the intramolecular channel of *Mt* ACS/CODH^[22]^. However, it is not known yet if the two branches are functional. The properties of the substrate channel(s) are likely to have a decisive impact on different catalytic properties of the enzyme, including the Michaelis constants for CO and for CO_2_ and the catalytic bias, as observed before with e.g. hydrogenases^[23,24]^. Additionally, since inhibitors such as O_2_ (whose physical properties are very similar to CO), cyanide^[25]^ or *n*-butyl-isocyanide^[21]^ most probably access the active site the same way as CO, the channel properties also modulate the strength of the inhibitors. For instance, it was shown that, for NiFe hydrogenases, CO and O_2_ (which are both inhibitors in this case) reach the active site at the same rate and using the same pathway, though O_2_ reacts more slowly with the active site than does CO^[24]^. Figure 1 shows parts of a sequence alignment of most of the CODHs (and ACS/CODHs) functionally characterized so far, focusing on the regions around the active site that contain the residues involved in the putative gas channels.

**Figure 1.**
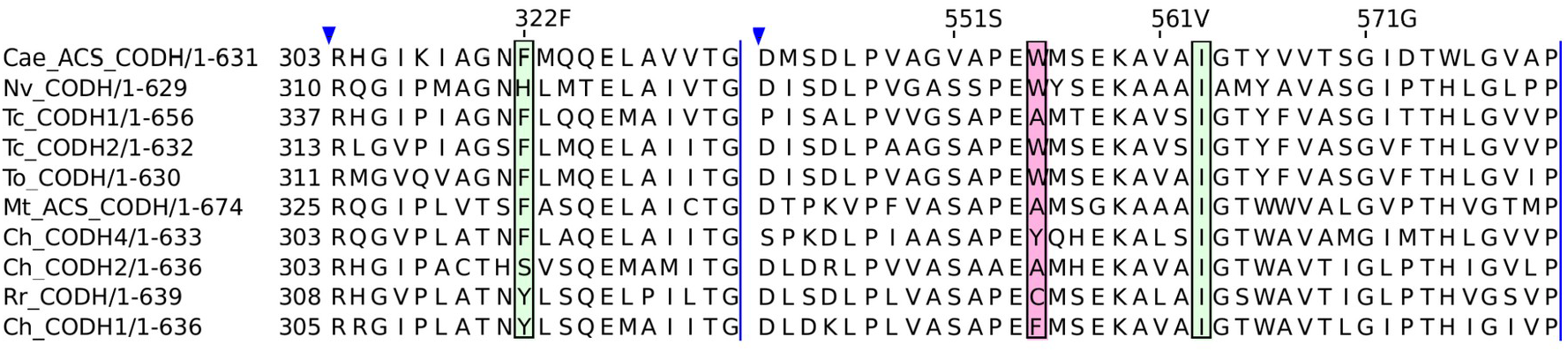
Sequence alignment of most of the CODHs (or ACS/CODHs) studied so far, emphasizing the two regions in which the variants have been constructed in this work. The green shaded regions correspond to the two residues mutated here (F322 and I563). The pink shaded region corresponds to the A559 residue in *Ch* CODH2 (corresponding to residue W555 in *Tc* CODH2). The sequences are those of *Clostridium autoethanogenum* ACS/CODH, *Nitratodesulfovibrio vulgaris* CODH, *Thermococcus sp AM4* CODH 1 and 2, *Thermococcus onnurineus* CODH, *Moorella thermoacetica* ACS/CODH, *Carboxydothermus hydrogenoformans* CODH4 and 2, *Rhodospirulum rubrum* and *Carboxydothermus hydrogenoformans* CODH1 (see the list of accession codes in SI, section S1). The numbering above the sequences is that of *Tc* CODH2.

Only three publications have addressed the effect of modifying the putative gas channel to the kinetic properties of CODHs (Michaelis constant and/or sensitivity to oxygen). Basak and coworkers recently described a series of site-directed variants of *Ch* CODH2, including mutations of the residue I567, which is conserved across all CODHs characterized so far^[26]^. The same residue, I560 in *Nv* CODH, is described as a Xenon contact site^[22]^, and in *Clostridium autoethanogenum* ACS/CODH, I564 lines the putative CO channel. Basak and coworkers found that mutating this isoleucine to leucine increases ten times the K_M_ for CO^[26]^.

Kim and coworkers conducted an extensive mutagenesis campaign to increase the oxygen resistance of *Ch* CODH2^[27]^. They formulated their working hypothesis on the observation that the A559 residue (*Ch* CODH2 numbering) is not conserved, and the size of the residue at this position seems to correlate with oxygen sensitivity: *Ch* CODH2 has an alanine, while *Ch* CODH4, which is much less sensitive than *Ch* CODH2, has a tyrosine, and *Nv* CODH, also not very sensitive, has a tryptophan. However, the comparison of the sequences actually suggests that the nature of the amino acid at that position is unlikely to be a general determinant of the resistance to O_2_: while *Ch* CODH4 has a tyrosine and resists the exposure to high O_2_ concentrations without inactivation, *Nv* CODH, which has a tryptophan, inactivates very quickly, but is resistant because it almost fully reactivates after an exposure to reducing conditions^[28]^. Furthermore, the two CODHs from *Thermococcus sp AM4* have a low resistance to O_2_ but, among the two, the most resistant to O_2_ is *Tc* CODH1, which has an alanine at that position, while *Tc* CODH2 has a tryptophan^[29]^. In the original article, Kim and coworkers produced a large number of mutants of the A559 residue and characterized their resistance to oxygen using solution assays; among other results, they claimed a one hundredfold increase in the resistance against oxygen when introducing a tryptophan^[27]^. In a follow-up article, they combined the A559W mutation with modifications of a surface residue, valine 610, to obtain even more O_2_-resistant variants^[30]^. However, when we characterized the A559W and A559W/V610H variants of *Ch* CODH2 using electrochemical methods, we could detect no impact of the mutations on the resistance of the enzymes against oxygen^[31]^.

In this work, we used site-directed mutagenesis to produce variants of *Thermococcus sp. AM4* CODH2 to study its gas channel. We examined the role of phenylalanine 322, a residue on the “backside” of the C-cluster, that was proposed to be a determinant of the high O_2_ resistance of *Ch* CODH4^[5]^, but we rule out its importance. We particularly focused on isoleucine 563, equivalent to I567 in *Ch* CODH2. We show that mutating this residue has important effects on the catalytic efficiency (100-fold reduction), K_M_ for CO, the catalytic bias and the resistance to O_2_. From our data, we conclude that size is not the dominant factor in slowing down the access to the active site, and that the rate of intramolecular diffusion of O_2_ and CO are proportional.

## Materials and methods

### Strains and plasmids

*E. coli* strain DH5α (F-, endA1, hsdR17(rK-mK +), supE44, thi-1, λ-, recA1, gyrA96, relA1, Δ(argF-lacZYA)U169, φ80dlacZΔM15) was used as a host for the construction of recombinant plasmids. The cultures were routinely grown at 37 °C in Luria–Bertani (LB) medium containing 20 μg/mL gentamicin when needed. *Solidesulfovibrio fructosivorans* (formerly *Desulfovibrio fructosovorans)* (*Sf*) strain MR400 (hyn::npt ΔhynABC) carrying a deletion in the [NiFe] hydrogenase operon^[32]^ was grown anaerobically for 5 days at 37 °C in fructose/sulfate medium as previously described^[33]^. 50 μg/mL kanamycin was present routinely, and 20 μg/mL gentamicin was added only when cells harbored the expression vectors.

The plasmid for the production of *Tc* CODH2 WT was constructed as described in ^[29]^. Site directed mutagenesis was performed on the *Tc* CODH2 WT plasmid using primers carrying the desired nucleotide substitutions. The primers are shown in supplementary Table S1. The entire plasmid was amplified by PCR with Pfu ultra HF from Agilent (600380). Following amplification, the PCR products were treated with DpnI from NEB (R0176S) to digest the parental methylated plasmid DNA. The resulting nicked circular mutant plasmids were transformed into *E. coli DH5α* competent cells from NEB (C2987H). Positive clones were screened, verified by Sanger sequencing, and introduced into the *Sf* strain MR 400 by electrotransformation. Exponentially growing cultures were harvested by centrifugation, washed several times with sterile cold water and resuspended at high density. Then mutant plasmids were introduced into the *Sf* strain MR400 by electroporation using Gene pulser apparatus (Bio-Rad) with optimized parameters for this strain (with 1 mm electroporation cuvettes: 1.5 kV, 25 µF, 400 ohms). Immediately after the electric pulse, cells were recovered in fructose sulfate medium, incubated under anaerobic conditions and, after 5 hours, the antibiotics were added to the cultures. After growth, 3 successive subcultures were subsequently carried out. The proteins were purified as described in ^[34]^ except that cells were disrupted in the glove box by either sonication or using the high pressure homogenizer Emulsiflex (Avestin Inc).

### Biochemical characterization

The CO oxidation activity of *Tc* CODH2 (WT and mutants) was measured in a glove box (Jacomex, filled with N_2_, O_2_ < 4 ppm) using a Varian Cary 50 spectrophotometer with a probe of 1 cm optical length. The CO oxidation activity was monitored at 37 °C by following the reduction of methyl viologen (MV) over time at 604 nm (ε = 13.6 mM^−1^ cm^−1^) in a plastic cuvette. Before recording the CO oxidation activity, the enzyme was diluted to a final concentration of 0.44 μM in 0.1 M Tris-HCl buffer, pH 8. To start the reaction, 5 μL of the enzyme solution were injected in a magnetic-stirred 1 mL cuvette containing 0.1 M Tris-HCl buffer (pH 8), 2.4 mM MV, 10 μM NaDT and 25 μM CO (25 μL of a CO-saturated solution injected just before the addition of the enzyme).

The nickel and iron content of the samples was determined using an iCAP 6000 ICP-OES spectrophotometer (ThermoFisher Scientific).

### Electrochemical experiments

Electroactive films of *Tc* CODH2 (WT and mutants) were prepared by simply depositing 0.3 μL of enzyme stock solution onto a pyrolytic graphite edge rotating disc electrode (PGE-RDE, 2.5 mm diameter) connected to an Autolab PGSTAT128N potentiostat (Metrohm, The Netherlands). The working electrode was rotated using an OrigaTrod electrode rotator (Origalys, France). All experiments were carried out in a Jacomex glove box filled with N_2_ (O_2_ < 4 ppm). All potentials are quoted with respect to the standard hydrogen electrode. The data was analyzed using the open source software QSoas^[35]^.

All the materials and methods are described in more detail in Supporting Information.

## Results

### Purification and biochemical characterization of the variants

We constructed the plasmids for the expression of 10 variants of *Thermococcus sp. AM4* CODH2: eight in which isoleucine 563 was replaced with a glutamic acid (I563E), a phenylalanine (I563F), a glutamine (I563Q), a glycine (I563G), an alanine (I563A), an arginine (I563R), a leucine (I563L), and a tryptophan (I563W), and two in which phenylalanine 322 was replaced with a histidine (F322H) and a serine (F322S). We wanted to change the size of the amino acid side chain as well as its functional group, since our previous experience with NiFe hydrogenases showed that not only the size but also the charge and/or hydrophilicity of residues in the channel may affect the rate of intramolecular diffusion^[36]^. We successfully transformed strains of *Solidesulfovibrio fructosivorans* with these plasmids, expressed and purified the proteins using a previously published method^[29]^. We determined their specific activity and metal contents. The I563Q, I563A and I563L variants have activities similar to that of the wild type enzyme, whereas the I563F, I563E and I563W variants suffer a factor of 3 to 4 loss in activity (table 1). Almost all of these variants have a Fe/Ni ratio similar to the WT (and to the expected value of 10), suggesting that they are properly matured. Similarly, both the F322H and the F322S are expressed and matured correctly, and have more than 50% of the activity of the WT. We could purify the I563R, but its activity was very low. We could not purify a significant amount of the I563G variant, nor detect any activity.

**Table 1.** Characterization of the wild-type and variants of *Tc* CODH2. The CO oxidation rate at pH 8 was measured in solution assays, the Fe/Ni content was determined by ICP-OES, all other parameters were determined by protein film electrochemistry. The conditions are described in SI sections S1 and S2. ND is not determined. All experiments were performed at 25 °C, the protein film voltammetry (PFV) CO oxidation experiments were performed at pH 7 and -0.31 V vs SHE, the CO_2_ reduction experiments (K_M_ and product inhibition) at pH 6 and -0.66 V vs SHE.

| | CO oxidation activity ($\mu\text{mol min}^{-1} \text{mg}^{-1}$) | Fe/Ni content (expected = 10) | Bias CO/CO<sub>2</sub> | K<sub>M</sub> for CO ($\mu\text{M}$) | k<sub>eff</sub> for CO ($\text{s}^{-1} \cdot \mu\text{M}^{-1}$) | K<sub>M</sub> for CO<sub>2</sub> (mM) | K<sub>i</sub> of CO<sub>2</sub> reduction by CO ($\mu\text{M}$) |
| --- | --- | --- | --- | --- | --- | --- | --- |
| WT | 400 ± 150 | 11.4 | 2 ± 1 | 0.8 ± 0.3 | 600 ± 400 | ND (too high) | 12 ± 3 |
| I563E | 98 ± 5 | 11.6 | 6 ± 1 | 4.3 ± 0.7 | 26 ± 6 | ND (too high) | 20 ± 3 |
| I563F | 65 ± 45 | 12.2 | 30 ± 20 | 4 ± 1 | 20 ± 20 | ND (too high) | 80 ± 38 |
| I563Q | 200 ± 20 | 10.9 | 28 ± 3 | 8 ± 1 | 30 ± 6 | ND (too high) | 40 ± 20 |
| I563A | 140 ± 60 | 14.2 | 32 ± 7 | 7 ± 1 | 20 ± 10 | ND (too high) | 116 ± 7 |
| I563W | 40 ± 20 | 10.2 | 50 ± 10 | 38 ± 11 | 1 ± 1 | ND (too high) | 150 ± 30 |
| I563L | 200 ± 150 | 8.7 | 3 ± 1 | 4 ± 1 | 60 ± 60 | ND (too high) | 10 ± 3 |
| I563G | ND | ND | ND | ND | ND | ND | ND |
| I563R | 3 ± 3 | 11.5 | ND | ND | ND | ND | ND |
| F322H | 150 ± 5 | 10.6 | 2.8 ± 0.2 | 0.9 ± 0.1 | 190 ± 30 | 1.5 ± 0.1 | 5 ± 1 |
| F322S | 110 ± 10 | 11.1 | 1.5 ± 0.2 | 0.5 ± 0.2 | 250 ± 100 | 2.9 ± 0.7 | 7 ± 1 |

### Electrochemical characterization of the variants: Michaelis and inhibition constants

To further characterize the variants and measure the Michaelis and product inhibition constants, we used the electrochemical approach^[37–39]^ previously developed for the characterization of the wild-type CODHs from *Thermococcus sp AM4*^*[29]*^.

Figure 2 shows experiments used to determine the value of the K_M_ for CO. The enzyme is immobilized onto an electrode that is immersed in the buffered solution of an electrochemical cell and poised at a high enough potential to drive the catalytic oxidation of CO. An aliquot of CO-saturated buffer is injected into the electrochemical cell solution, yielding an instant increase in the dissolved CO concentration, followed by a slow exponential decrease that results from the exchange of CO between the cell solution and the CO-free atmosphere of the glove box^[40]^. The time-dependent CO concentration is shown in figure 2A. Figure 2B shows the response in current of a film of *Tc* CODH2 WT. A catalytic oxidation current appears instantly after the injection of CO. It remains nearly constant for a while, despite the decrease in CO concentration, because the latter is still above the K_M_ value. Then, the current decreases, ending in a perfect exponential decay, when the CO concentration is below K_M_ and the catalytic rate is proportional to CO concentration. An appropriate modeling of the data, taking into account the mass-transport kinetics of CO towards the electrode^[41]^, allows the determination of the K_M_ value (here 0.8 µM); the perfect fit of the model is shown as a dashed red trace. Figure 2C shows the response of the I563E variant of *Tc* CODH2 to an identical injection of CO. The initial period where the current is nearly constant is significantly shorter than in the case of the WT, meaning that the K_M_ for CO of this variant is greater than that of the WT: indeed, the modeling gives a value of 3.3 µM. The results for all the enzyme variants studied in this work are shown in table 1. They are the average of several duplicates (at least 3). The most striking effect is the 50-fold increase in the value of K_M_ that results from the I to W mutation.

**Figure 2.**
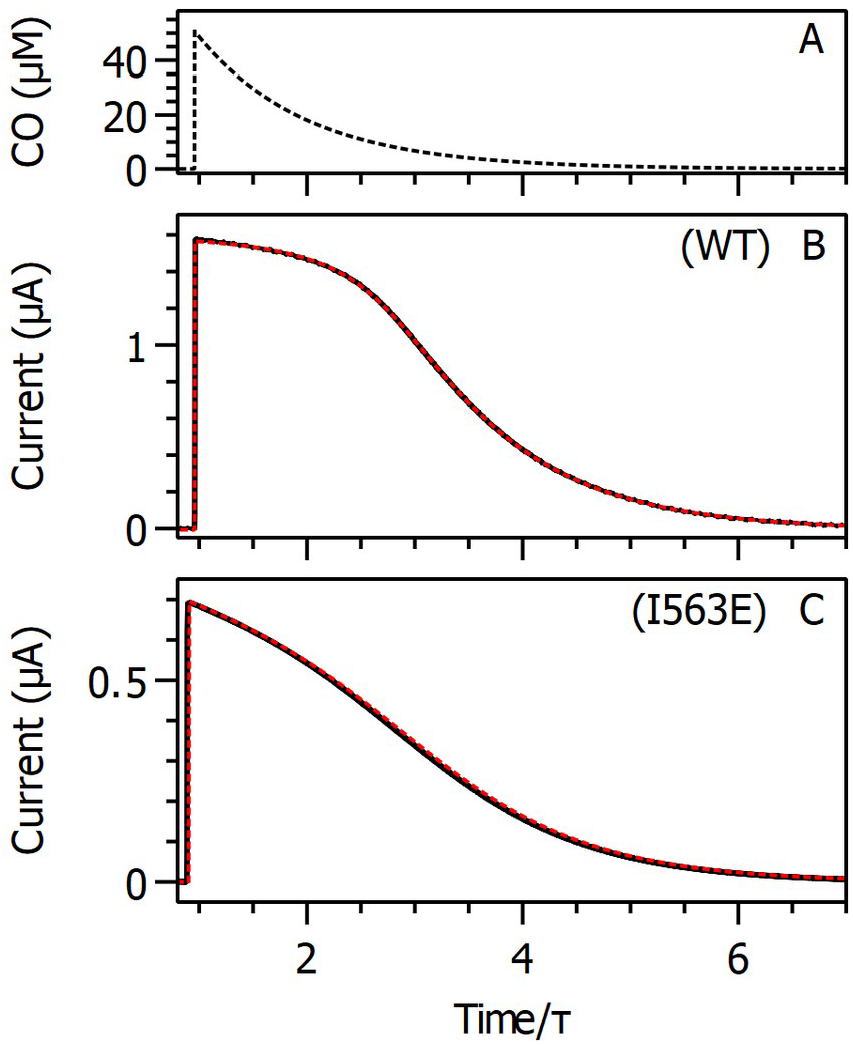
Chronoamperometric measurements of the Michaelis constant relative to CO. On the x axes, the time is normalized by the characteristic time of departure of CO from solution (determined from the fits of Eq. (9) (model c) in ^[41]^ to the data). A) CO concentration in the cell vs. time. B and C) Black lines: current observed with films of *Tc* CODH2 WT (B) and I563E variant (C), submitted to a single injection of 50 μM CO at t = 0 (the CO departure time constants are τ = 42 s and 44 s for B and C, respectively). Experimental conditions: T = 25 °C, pH 7, electrode rotation rate ω = 4000 rpm, E = −0.31 V vs. SHE. Red dashed lines: fit of the chronoamperograms using Eq. (9) (model c) in ^[41]^. Parameters of the fits: B) K_M_ = 0.96 μM, nFAμ = 1.68 μA/μM, nFAm = 0.21 μA/μM, i_0_ = −6.4 nA, [CO]_0_ = 50 μM (not adjusted), τ = 42 s, and C) K_M_ = 3.3 μM, nFAμ = 0.25 μA/μM, nFAm = 0.17 μA/μM, i_0_ = −0.19 nA, [CO]_0_ = 50 μM (not adjusted), τ = 44 s. A is the electrode area, F the Faraday constant, n (=2) the number of electrons. m is the mass-transport coefficient of CO, and µ the product of the electroactive coverage and the enzymatic catalytic efficiency (see details in ref. ^[41]^).

The same experiment can be used in principle to determine the value of the K_M_ for CO_2_. However, for all but the F322 mutants, the value of the K_M_ was larger than the highest concentration of CO_2_ we could inject without changing the pH too much (about 7 mM), thus being impossible to determine. The reduction of CO_2_ is strongly inhibited by the presence of the product CO. Like before^[29]^, we have used injections of CO_2_ followed by injections of CO to determine it for all the variants. The inhibition constant for most variants is so small that the enzyme is inhibited to some extent by the CO produced by the reaction and not yet spun away by the electrode rotation; the data analysis takes this into consideration (see details in SI section S2).

### Determination of the catalytic bias

The catalytic bias can be defined as the ratio between the catalytic rates of CO oxidation and CO_2_ reduction measured under very oxidizing or reducing conditions, respectively, and in the absence of the substrate for the other direction^[38]^. As the catalytic rates for the two reactions are determined under completely different conditions, under which it is expected that the reaction is unidirectional (only one substrate, high driving force), their ratio is not defined by thermodynamics, but rather represents an intrinsic property of the enzyme^[23,42]^.

We used cyclic voltammetry to determine the value of the catalytic bias for the WT and the variants. Figure 3 shows cyclic voltammograms recorded in the presence of either CO or CO_2_, for all of the active CODHs described in this work. As the electroactive coverage can vary from one film to another, for each variant the two CVs were recorded with the same film, allowing the quantitative comparison of the oxidation and reduction currents (additional precautions were used in order to minimize the impact of the decrease of electroactive coverage over time, see SI section S2). We used the same experimental conditions as in our previous work^[29]^. For *Tc* CODH2 WT, the magnitudes of the CO oxidation current and the CO_2_ reduction current are comparable, showing that this enzyme is not particularly biased in either direction. This is also the case of the two F322 variants. However, mutating the I563 residue has a strong impact on the bias, decreasing the CO_2_ reduction current much more than the CO oxidation current, resulting in a shift of the bias towards CO oxidation. The numerical values of the bias reported in table 1 were determined from the ratios of the oxidation current at −160 mV in the presence of only CO and the reduction current at −760 mV in the presence of only CO_2_.

**Figure 3.**
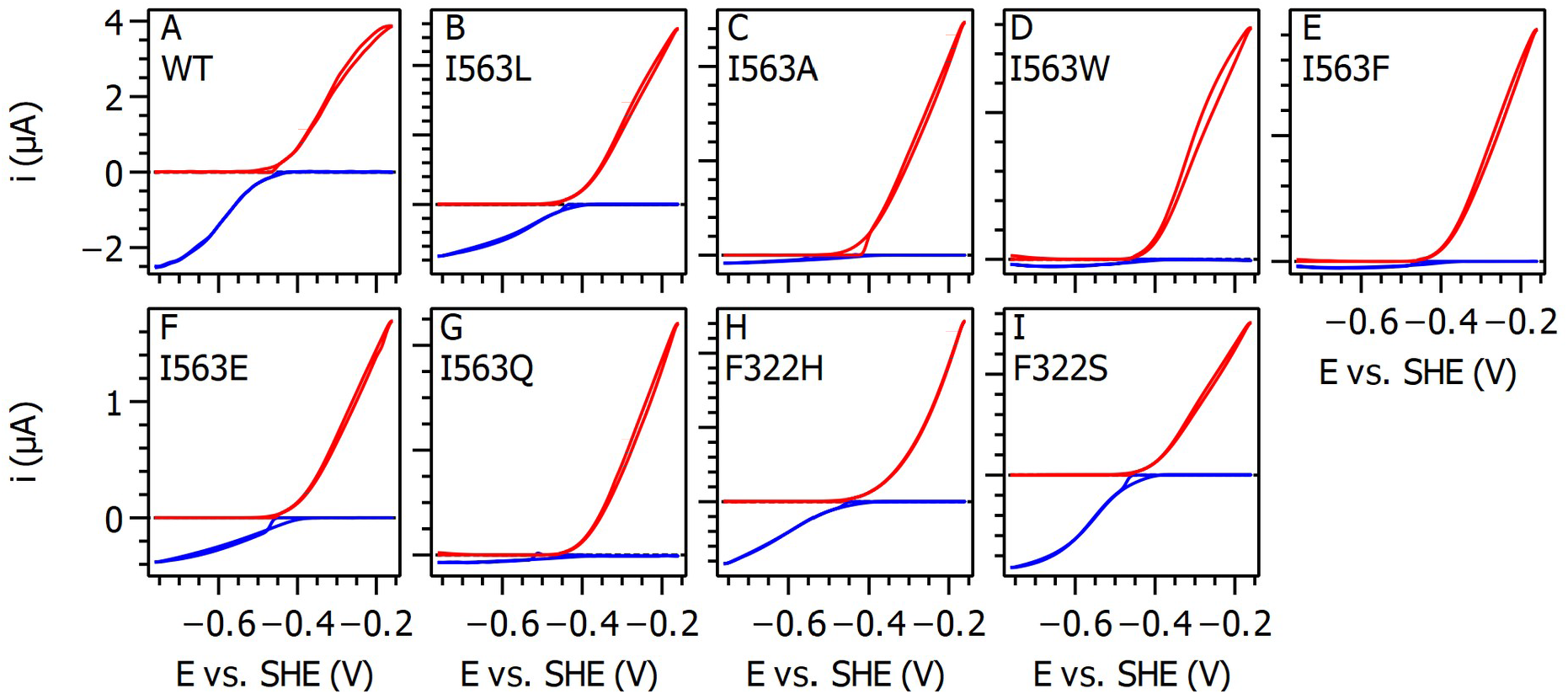
Dependence of CO oxidation and CO_2_ reduction activities as a function of the potential for all the active enzymes described in this article (the enzymes are indicated directly on the panels). The red CVs were recorded in the presence of 50 µM CO (and no CO_2_), the blue CVs in the presence of 7 mM CO_2_ (and no CO). In all the cases, the same enzyme film was used to record both CVs. All the CVs were baseline-subtracted to ease the comparison. Conditions: T = 25 °C, pH 6 (0.2 M MES), ω = 4000 rpm, scan rate = 20 mV/s.

### Reactivity with O_2_

Last, we characterized the reactivity of all the variants with oxygen using a method that we developed earlier^[28]^. This method consists in poising the electrode at a constant potential and exposing a film of CODH to six injections of CO to detect the CO oxidation activity, one injection of O_2_ (shortly after the third CO injection) to inhibit the enzyme, and a low potential reductive poise to trigger reductive reactivation. All together, the data can be used to quantitatively assess the effect of exposure to O_2_ and reductive activation, with little sensitivity to film loss.

The method is illustrated in figure 4, which shows the concentration of CO and O_2_ in panel A, and the responses in current of a film of WT (panel B) and a film of the I563F variant (panel C). Injections 1 and 2 (around t = 40 s and t = 400 s) are similar to those shown in figure 2, with a small decrease in current that can be attributed to film loss. Injection 3 (around t = 820 s) is followed by the injection of O_2_ resulting in a fast decrease of the current, and a slow recovery (almost invisible in the case of the WT). Subsequent injections (4, 5 and 6) show some CO oxidation activity, although significantly less than after injections 1 and 2. A reductive poise precedes injection 5, which makes it possible to assess the extent of reductive reactivation. The data in figure 4 also shows that the activity remaining after exposure to O_2_ is much greater in the case of the I563F variant than in the case of the WT, showing that the former is more resistant to O_2_ than the latter.

**Figure 4.**
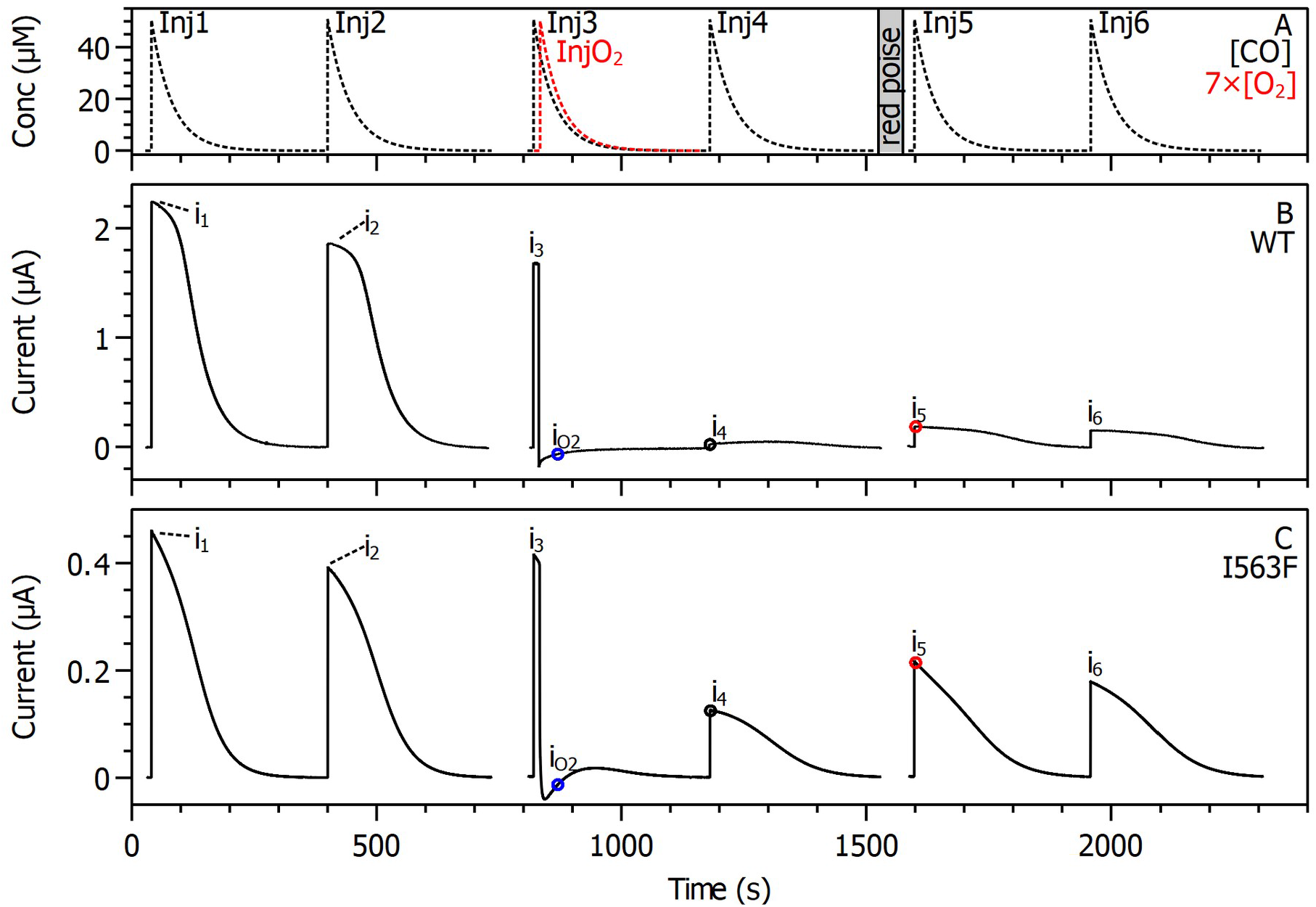
The 7-injection experiments used to characterize the response of CODHs to exposure to O_2_. Panel A shows the concentration of CO (black) and O_2_ (red) over time, showing their exponential decrease in solution after each injection (6 injections for CO, and 1 injection for O_2_). Panels B and C show the responses in current of a film of WT (B) and of the I563F variant (C).

Fitting an exponential decay to the fast decrease in current immediately after the O_2_ injection allows the determination of *k*_*i*_, the pseudo-first order rate constant of inhibition by O_2_. Comparing the currents just after the injection of O_2_ (i_O2_) and injections 4 and 5 (i_4_ and i_5_) to that just after injection 3 (i_3_) allows the determination of the fractions of activity remaining immediately after O_2_ injection, activity recovered after O_2_ departure, and activity recovered after a low-potential poise (at -660 mV). In practice, all six CO injections are necessary to discriminate between the effects of O_2_ inhibition and film loss; the details are explained in ref ^[28]^ and SI section S3. This series of 7 injections was repeated by systematically varying the concentration of O_2_ for each enzyme variant; the results are shown in figure 5 and figure 6.

**Figure 5.**
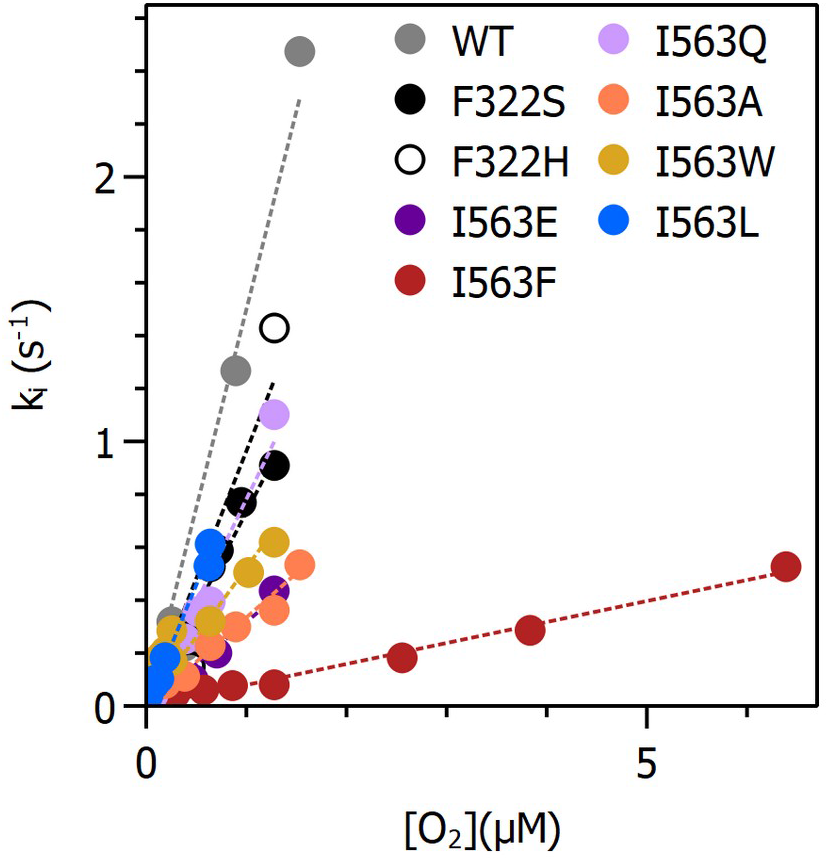
Pseudo-first order rate constants of inhibition of *Tc* CODH2 wild-type and variants as a function of injected O_2_ concentration. The color code is indicated on the graph. The lines are the linear fits, which give the second-order rate constants shown in table 2. Conditions: T = 25 °C, pH 7, ω = 4000 rpm, E = −0.31 V vs. SHE.

**Figure 6.**
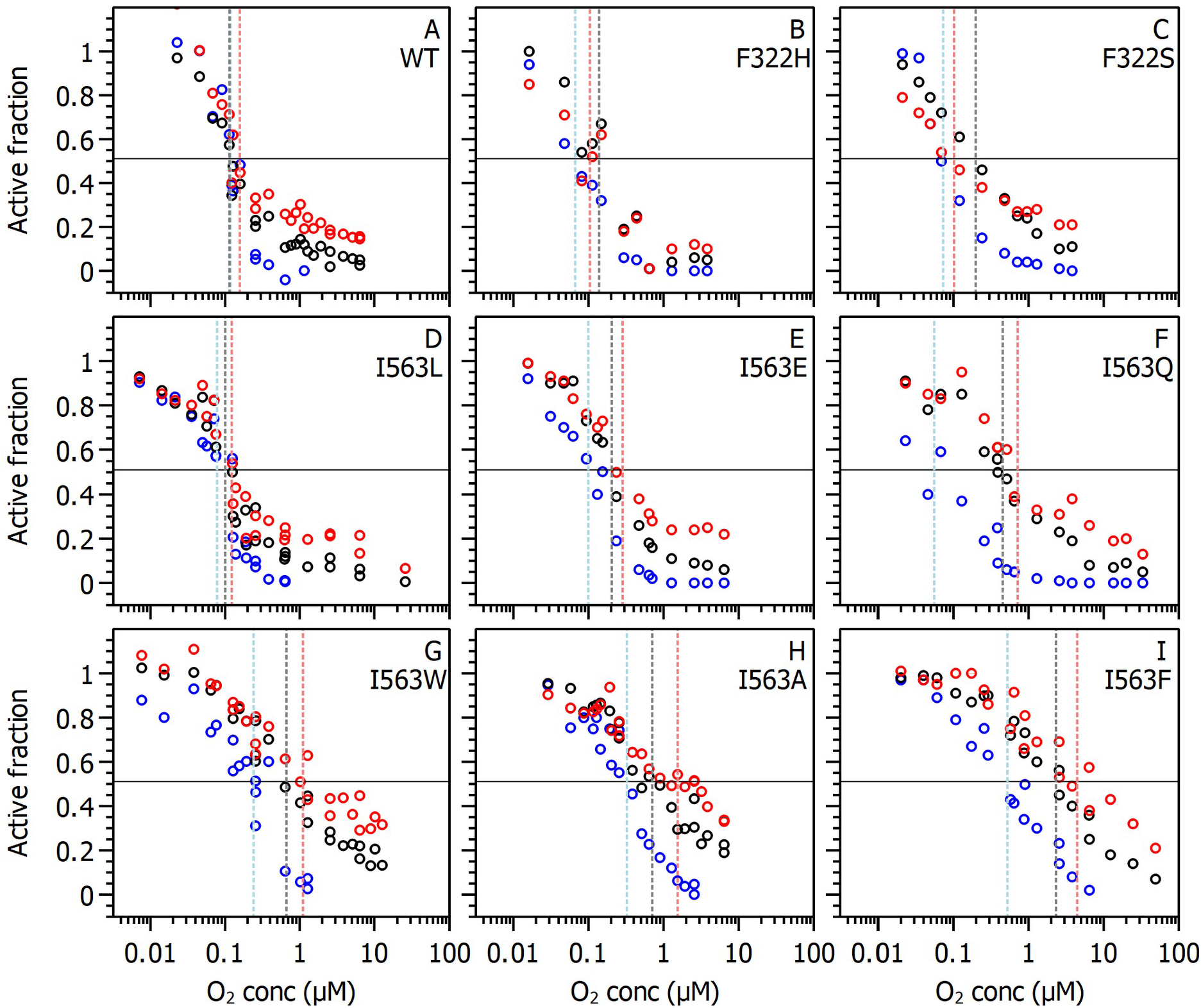
Effect of different O_2_ concentrations on all the enzymes studied in this work, as deduced from the 7-injections experiments in PFV (the variants are indicated directly on the panels). Blue circles: activity remaining just after O_2_ injection; black circles: activity recovered after O_2_ removal; red circles: activity recovered after a poise at low potential (E = −0.66 V vs. SHE for 10 s). The vertical lines indicate the resulting 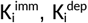 and 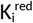 values (see SI section S3). Conditions: T = 25 °C, pH 7 (mixed buffer), ω = 4000 rpm, E = −0.31 V vs. SHE. Supplementary figure S4 shows the same data, but grouped by 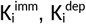 and 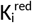 rather than by variant.

Figure 5 shows the pseudo-first order rate constants of inhibition by O_2_ as a function of O_2_ concentration for all the CODHs characterized in this work. In all cases, the pseudo-first order rate constants are proportional to the concentration of injected O_2_, which made it possible to use the slope of each line to determine the bimolecular rate constant of reaction with O_2_, *k*_*O2*_ (Table 2).

**Table 2.** Parameters that quantify the inhibition by O_2_ of *Tc* CODH2 and its variants. The first three parameters represent the concentration of O_2_ resulting in 50% of the activity being lost: just after O_2_ injection 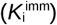, after O_2_ departure 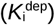, and after a reductive poise 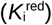 . The fourth parameter is the bimolecular rate constant of reaction with O_2_, *k*_*O2*_. Conditions: T = 25 °C, pH 7, ω = 4000 rpm, E = −0.31 V vs. SHE. *k*_*corr*_ is the value of the bimolecular rate constant, corrected under the assumption that the inhibition by O_2_ is competitive: *k*_*corr*_ *= k*_*o2*_ × (1+ [CO]/ *K*_*M*_), in which [CO] is the concentration of CO at the injection of O_2_, here 50 µM.

| Variant | $K_i^{\text{imm}}$ ($\mu\text{M O}_2$) | $K_i^{\text{dep}}$ ($\mu\text{M O}_2$) | $K_i^{\text{red}}$ ($\mu\text{M O}_2$) | $k_{O_2}$ ($\text{s}^{-1} \mu\text{M}^{-1}$) | $k_{\text{corr}}$ ($\text{s}^{-1} \mu\text{M}^{-1}$) |
| --- | --- | --- | --- | --- | --- |
| WT | $0.12 \pm 0.01$ | $0.114 \pm 0.009$ | $0.16 \pm 0.01$ | $1.5 \pm 0.2$ | $100 \pm 50$ |
| I563E | $0.0992 \pm 0.008$ | $0.20 \pm 0.01$ | $0.28 \pm 0.02$ | $0.32 \pm 0.04$ | $4 \pm 1$ |
| I563F | $0.52 \pm 0.02$ | $2.3 \pm 0.2$ | $4.4 \pm 0.9$ | $0.079 \pm 0.005$ | $1.1 \pm 0.3$ |
| I563Q | $0.06 \pm 0.01$ | $0.452 \pm 0.009$ | $0.72 \pm 0.06$ | $0.78 \pm 0.09$ | $6 \pm 1$ |
| I563L | $0.077 \pm 0.005$ | $0.100 \pm 0.004$ | $0.123 \pm 0.005$ | $0.91 \pm 0.07$ | $12 \pm 4$ |
| I563A | $0.324 \pm 0.007$ | $0.71 \pm 0.05$ | $1.5 \pm 0.1$ | $0.33 \pm 0.06$ | $2.7 \pm 0.9$ |
| I563W | $0.24 \pm 0.01$ | $0.66 \pm 0.04$ | $1.11 \pm 0.05$ | $0.5 \pm 0.1$ | $1.2 \pm 0.6$ |
| F322H | $0.0665 \pm$ | $0.14 \pm 0.02$ | $0.10 \pm 0.02$ | $1.0 \pm 0.2$ | $60 \pm 20$ |
|  | 0.002 |  |  |  |  |
| F322S | $0.073 \pm 0.004$ | $0.197 \pm 0.003$ | $0.102 \pm 0.005$ | $0.73 \pm 0.09$ | $70 \pm 40$ |

Figure 6 shows the remaining activity just after O_2_ exposure (in blue), after O_2_ departure (in black) and after a reductive poise (in red) as a function of the injected concentration of O_2_ for all the variants. We re-run the characterization of *Tc* CODH2 WT^[29]^.

All the variants show a similar behaviour: of course, the activity is almost fully retained after injections of very small O_2_ concentrations, and the fraction of remaining activity decreases upon increasing the O_2_ concentration. In all cases, the activity remaining just after the injection is smaller than that left after O_2_ departure, itself smaller than that after reduction especially at higher oxygen concentrations, showing that the enzyme partially reactivates when O_2_ is gone, and by reduction. As was remarked in the first work of this kind on *Nv* CODH and *Ch* CODH2, this observation implies that the inactivation reaction produces at least three species, one that reactivates immediately, one that reactivates upon reduction, and one that cannot be reactivated^[28]^.

The impact of the mutations can be seen by the fact that the features of the curves in figure 6 significantly change from one variant to the other: the O_2_ concentrations that are necessary to decrease the activity, how much activity is recovered by removing O_2_ or reducing the enzyme. For instance, in the case of the WT (panel A), the three curves are almost superimposed, showing that very little activity is recovered after the departure of O_2_ and by reduction of the enzyme. In the case of the I563F variant (panel I), the curves are all shifted to greater O_2_ concentrations, and the red and black curves are shifted to even greater O_2_ concentrations compared to the blue one, showing that this variant is more resistant to O_2_ than the WT, and that removing O_2_ and reducing the enzyme help recover further activity.

To compare the O_2_ sensitivity of the variants, like in reference ^[29]^, we extracted three “apparent inhibition constants” from each series of experiments, which correspond to the concentrations of O_2_ that yield 50% remaining activity: 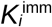 corresponds to 50% activity remaining just after O_2_ exposure, 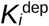 to 50% activity remaining after O_2_ departure, and 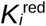 to 50% activity remaining after the reduction; these values are akin to IC50 values. All these apparent inhibition constants are reported in table 2, and also shown in figure 6 as vertical dashed lines. These values are only “apparent inhibition constants”, not true inhibition constants, since the inhibition is not fully reversible.

From these results, the different *Tc* CODH2 variants studied in this work can be classified into several groups. The F322H, F322S and I563L variants have the same behaviour as the WT, with only small variations in the values of the *K*_i_ and no significant changes in the bimolecular rate constant of reaction with O_2_. The I563Q mutation slightly increases the values of *K*_i_. The I563E variant has similar values of *K*_i_ as the WT in spite of a fivefold decrease in the bimolecular rate constant of reaction with O_2_. In the case of the I563A and I563W variants, the values of the *K*_i_ are significantly greater than those of the WT, and the bimolecular rate constant is three to five times smaller. The I563F variant features both a 20-fold decrease in the bimolecular rate constant and a 20 to 30-fold increase of both 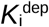 and 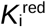.

### In-silico characterization of the channels

We used the software CAVER^[43]^ to determine the possible channels linking the buried active site to the solvent (see SI section S5 for more information). The network of channels determined this way is shown in figure 7A, with a zoom around the active site in figure 7C. The computations show an initial narrow channel that branches at 9 Å from the active site, precisely at the position occupied by I563, as described before^[22,27]^. Here, these branches will be referred to by the name of one of the residues lining them, either F567 or V561. The channels subdivide further along the path to the solvent, but these regions are most likely too far from I563 to be affected by the point mutations; we did not consider them further.

**Figure 7.**
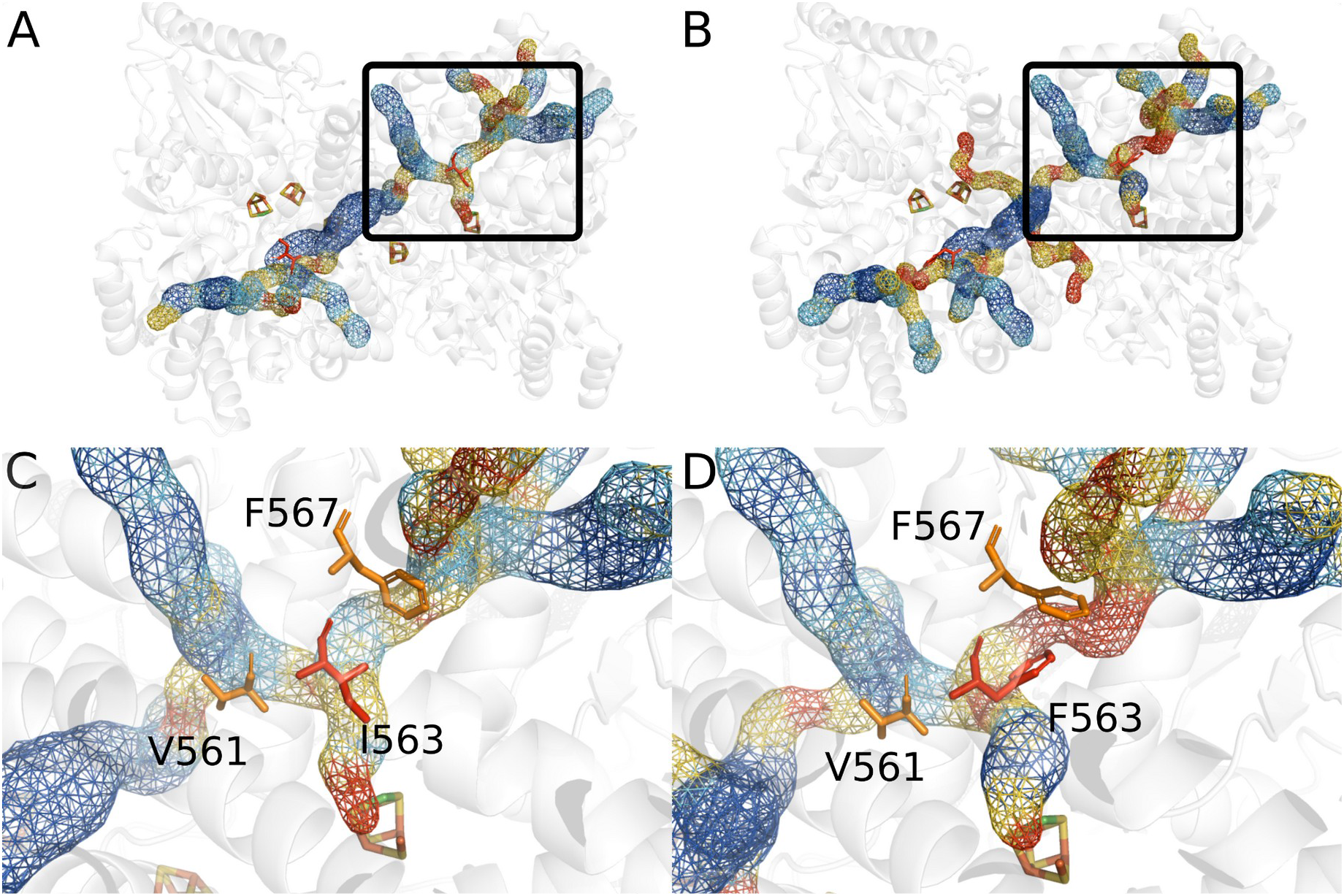
Panels A and C: *Tc* CODH2 WT crystal structure (PDB 6T7J) with the computed tunnels. Panels B and D: an AlphaFold model of the *Tc* CODH2 I563F variant, with the tunnels computed with CAVER. In panels C and D, the residues V561 and F567 are represented in sticks and colored in orange to indicate the 2 branches of the tunnel after the branching point. The residue in position 563 is highlighted in red sticks. Color code for the tunnel radius: red from 0.9 to 1.3 Å, yellow from 1.3 to 1.7 Å, light blue from 1.7 to 2 Å, and blue if above 2 Å. For the sake of the discussion, we have used the fully matured active site of *Ch* CODH2 (PDB: 3B51) to better visualize the position of the Ni atom and gauge the orientation (the active site in structure 6T7J is devoid of nickel).

We systematically used AlphaFold^[44]^ to determine the structure of each of the I563 variants, and CAVER to examine the influence of the mutations on the geometry of the channels. Figure 7B and D show the results for the I563F variant. The overall layout of the channels does not change (with the exception of the I563W variant, in which the F567 branch is closed). However, the channel width is affected by the mutations. For instance, in the case of the I563F variant, figure 7D shows that one of the channels is significantly constricted compared to the WT (the red region, located around residue F567), while the common part near the active site is actually enlarged, featuring a new cavity.

We have characterized the width profile along the two tunnels for the WT and all of the variants (details in supplementary section S5). We report in table 3 the width of the narrowest element of the two channels for each variant.

**Table 3.**
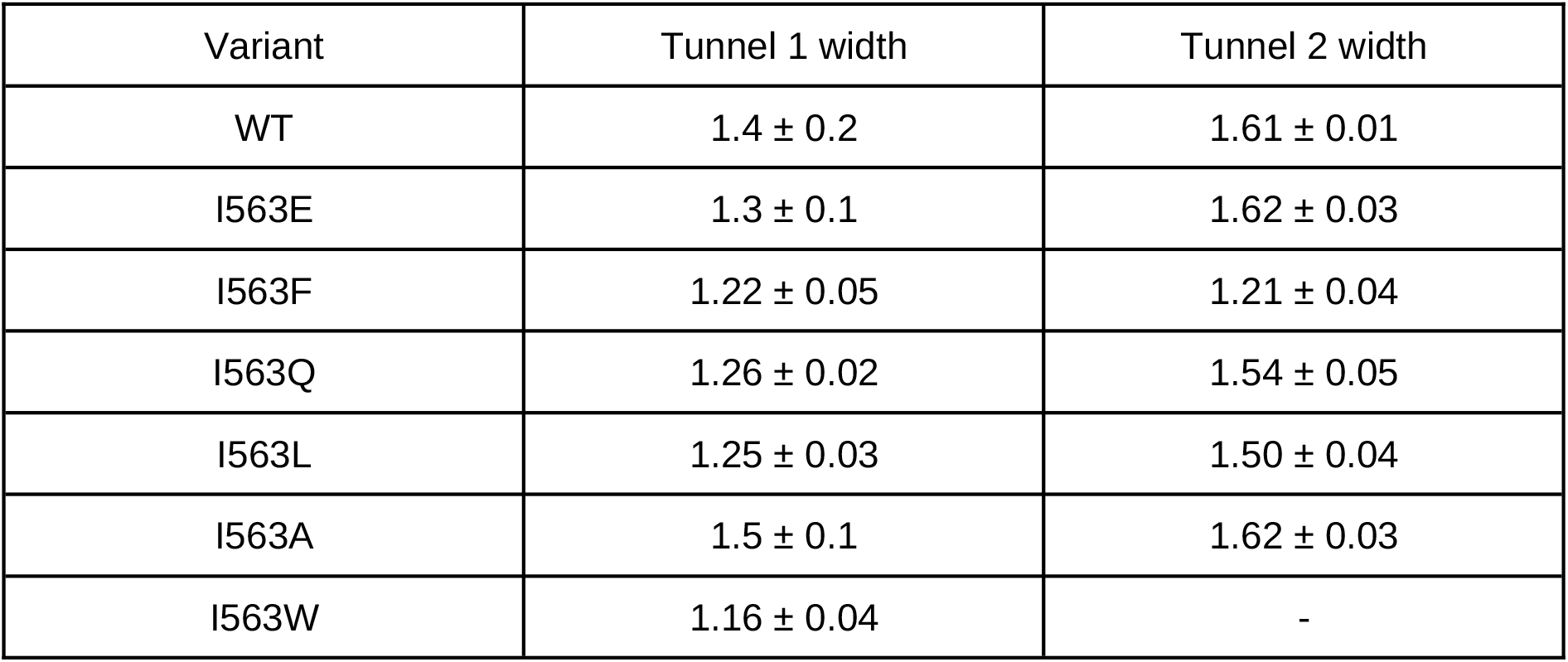
The width of the narrowest position of the tunnels determined using AlphaFold and CAVER. The error bar corresponds to the average over 5 different structures proposed by AlphaFold. In tunnel 2 of I563W no width is provided as the second channel is closed in this variant.

## Discussion

We have studied two series of variants of *Thermococcus* CODH2 obtained by replacing residues F322 and I563. The F322 residue is equivalent to F312 of *Carboxydothermus hydrogenoformans* CODH4, and it was initially proposed to play a role in the oxygen resistance of this enzyme^[5]^, since the only difference between *Ch* CODH2 and CODH4 around the active site is that residue: it is a serine in *Ch* CODH2. However, we have shown here that, in the case of *Thermococcus sp. AM4*, replacing this residue with histidine or serine has little impact on O_2_ resistance or on the biochemical parameters of the enzyme. The exceptional O_2_ resistance of *Ch* CODH4 must therefore find its origin in other residues, further away from the active site.

Concerning position 563, tables 1 and 2 show that the mutations impact both the reactivity with CO/CO_2_ and O_2_. However, with the exception of the I563R substitution, the specific activities are only moderately affected, suggesting that the mutations impact the transport towards the active site rather than the active site chemistry. These results are in line with those of Basak and coworkers, who found that point mutations of I567 in *Carboxydothermus hydrogenoformans* CODH2 change the Michaelis constant for CO but barely affect the enzymatic activity^[26]^. The impact of the I563 mutations on the catalytic efficiency is important, ranging from a factor of 10 (I563L) to more than two orders of magnitude for the I563W variant. This confirms that the I563 residue does line the gas channel, and that the mutations slow down the access of CO to the active site. Consistently with the 10-fold increase in the K_M_ observed by Basak and coworkers for the I567L variant of *Ch* CODH2^[26]^, even the conservative isoleucine-to-leucine mutation significantly impairs the catalytic efficiency, showing that the isoleucine is highly tuned, as can be expected from its very high conservation. The size of the amino acid introduced at position 563 is not the main criteria for the catalytic efficiency: if the I563W variant is indeed the slower variant, there is virtually no difference between the alanine and phenylalanine variants.

Figure 8 shows the catalytic efficiency as a function of the average of the width of the narrowest element of both channels, as deduced using AlphaFold and CAVER. The catalytic efficiency decreases strongly as a result of narrowing the channels, with the exceptions of the alanine and glutamate variants, which are predicted to have openings similar to the WT, but nevertheless a significantly smaller catalytic efficiency. It may be that their structures or channels are badly predicted, or that the correlation breaks down because these variants can accommodate a water molecule in the channel.

**Figure 8.**
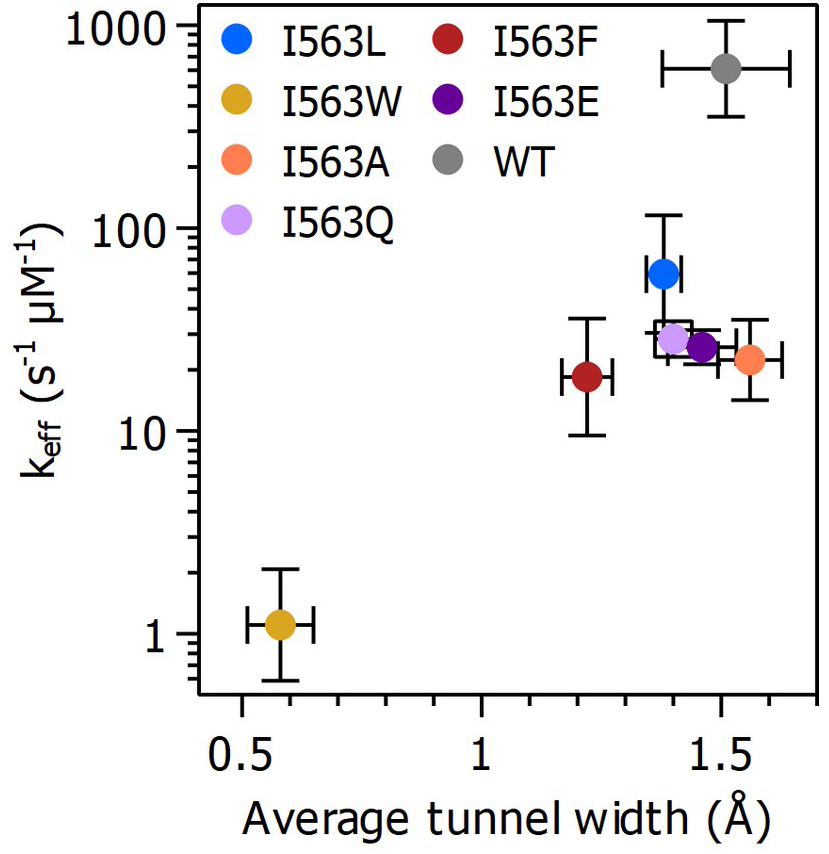
Plot of catalytic efficiency as a function of the average of the narrowest path for tunnels 1 and 2, and predicted from AlphaFold and CAVER.

Table 1 shows that all the variants have an increased bias for CO oxidation over CO_2_ reduction compared to the wild type. This implies that the constriction of the channel impacts the transport of CO_2_ to the active site more than that of CO. Indeed, figure 9A shows that the bias scales as the reciprocal of the catalytic efficiency, which suggest that the impact of the mutation on CO_2_ reduction is roughly the square of the impact on CO oxidation.

**Figure 9.**
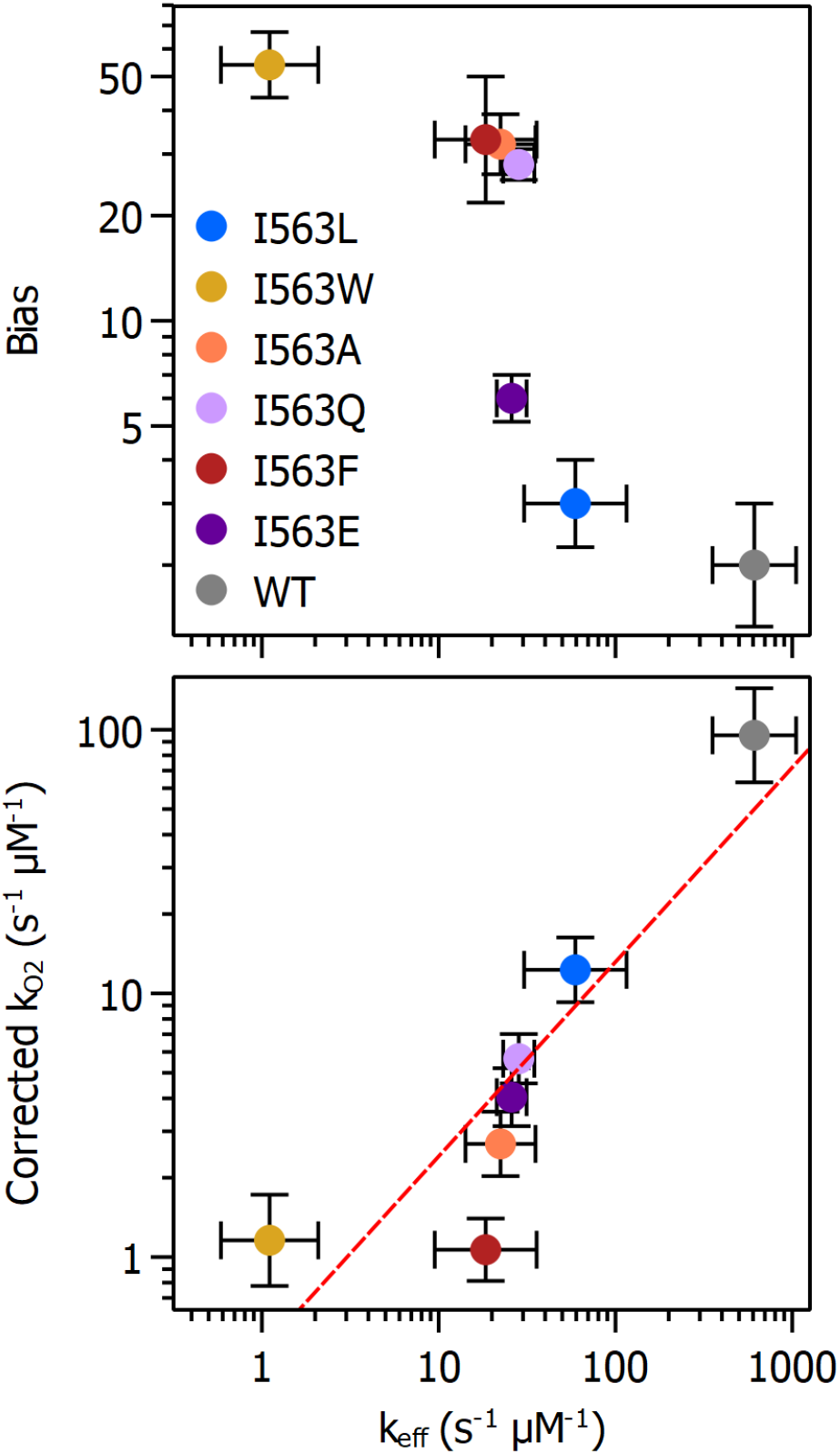
Plot of the catalytic bias (panel A) and the corrected bimolecular rate constant of reaction with oxygen (panel B) as a function of the catalytic efficiency. The data come from tables 1 and 2; the corrected bimolecular rate constant is computed thus: 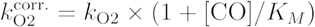, in which the concentration of CO is the one at the injection of O_2_, i.e., 50 µM. To avoid cluttering the top panel, the names of the variants are only given on the bottom. The red line in panel B has slope unity, showing that the rates of reaction with O_2_ and CO are proportional to one another.

Concerning the reactivity with O_2_, there is no obvious pattern in the raw data of table 2. However, it is possible to observe a clear trend if one assumes that the inhibition by O_2_ is competitive. This is a reasonable assumption because O_2_ targets the active site^[45]^, and the competitive inhibitor n-butyl-isocyanide protects from O_2_ damage^[28]^. The corrected value of the bimolecular rate constant of reaction with O_2_ shows an almost perfect linear correlation with the catalytic efficiency (the ratio k_cat_/K_M_, which is the bimolecular rate of reaction of the enzyme with CO, figure 9B). Applying a similar correction yields similar correlations for the IC50 values (SI figure S3). These results suggest that O_2_ is indeed a competitive inhibitor, and that the rates of intramolecular diffusion of O_2_ and CO are proportional to each other: slowing down the ingress of O_2_ necessarily slows down that of CO. This conclusion contrasts with the claims of Kim and coworkers in their study of *Carboxydothermus hydrogenoformans* CODH2^[27,30]^. Our conclusion is in line with the extensive mutagenesis study of the gas channel of the NiFe hydrogenase from *Solidesulfovibrio fructosivorans* (formerly D. fructosovorans) by Liebgott and coworkers, who showed that in a large series of gas channel variants, the intramolecular rates of diffusion of the inhibitors CO and O_2_ are proportional to each other^[24]^.

In the experimental conditions that we used, corresponding to about 5% CO (50 µM), the I563F variant shows a remarkable increase in the resistance against O_2_, a 20 to 30-fold increase in 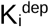 and 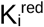 . This result is remarkable: with the exception of the claims by Kim and coworkers (which we could not reproduce^[31]^), this is the first report of a mutation of CODH that significantly protects the enzyme from O_2_ damage. The I563F variant strikes an optimal compromise in which O_2_ protection is significantly increased, but the penalty in terms of catalytic efficiency and K_M_ is reasonably small. However, our results demonstrate that the strategy that consists in blocking the gas channel cannot afford further protection without having a steep cost in terms of catalytic performance.

## Conclusion

Using protein engineering to increase the resistance of CODH to O_2_ is a prerequisite to any use of these enzymes for the reduction of CO_2_, but despite recent progress using crystallography^[45,46]^ and electrochemistry^[28,47]^, not much is known about the reaction of CODH with O_2_. In certain CODHs, the inhibition is partly reversible: some activity is recovered even under oxidizing conditions after anaerobicity is restored, and further reactivation can be induced by the reduction of the enzyme^[28,47]^. The latter reactivation results from the formation, upon exposure to O_2_, of a well-defined form of the active site that is inactive but protected from O_2_^[46]^. Like in the case of hydrogenase, the inhibition of CODH by O_2_ is therefore a complex process, and there is no unique parameter that would quantify an “overall sensitivity”. The inhibition depends on how fast the enzyme reacts with O_2_, but also on whether the product(s) of the reaction with O_2_ can be reactivated, how fast and under which conditions of potential. Understanding the mechanism of reaction with O_2_ and comparing the sensitivity of different enzymes and variants requires each of these aspects to be considered, which is possible when one uses direct electrochemistry experiments like those in figure 4.

Previous attempts to use protein engineering to increase the O_2_ resistance of CODH have led to controversial results^[31]^. Here we could demonstrate that the conserved I563 residue determines the rate of transport of both CO and O_2_. Its replacement simultaneously and significantly increases the K_M_ for CO (up to 40-times), decreases the catalytic efficiency (by almost 3 orders of magnitude), and increases the resistance to O_2_ (IC50s increased by up to 30 times). These effects are all strongly correlated because CO and O_2_ have similar sizes and use the same pathway, so that it is not possible to prevent O_2_ access to the active site without also decreasing the activity of the enzyme for its substrate. The comparison of the properties of the large series of CODH variants described here does not support the previous hypothesis in refs ^[27,30]^ that different pathways are used by CO and O_2_ and could be selectively blocked to protect the enzyme from O_2_ without any negative trade-off on catalytic activity.

## Acknowledgements

The authors acknowledge support from CNRS, Agence Nationale de la Recherche (grants ANR-17-CE11-0027, ANR-21-CE50-0041, ANR-23-SODR-0004, ANR-23-CE44-0046), and Region PACA. This project has received funding from the European Union’s Horizon Europe research and innovation programme under grant agreement number 101115403 (ECOMO) and from European Union HORIZON-TMA-MSCA-SE (grant agreement No. 101183014). This work has benefited from French State aid managed by the Agence Nationale de la Recherche under France 2030 plan, bearing the reference code ANR-22-PESP-0010: Projet ciblé “POWERCO2” within the PEPR project SPLEEN. The project leading to this publication has received funding from Excellence Initiative of Aix-Marseille University— A*Midex, a French “Investissements d’Avenir” program. The authors are members of the French Bioinorganic Chemistry group (http://frenchbic.cnrs.fr).

## Supporting informations

Supporting information contains all materials and methods, together with supplementary figures S1 to S3.

