## Supplementary information for "A conserved isoleucine gates the diffusion of small ligands to the active site of NiFe CO-dehydrogenase"

### - Supporting information -

Laura Opdam, Marta Meneghello+, Chloé Guendon, Jade Chargelègue, Andrea Fasano, Aurore Jacq-Bailly, Christophe Léger, Vincent Fourmond*

Aix-Marseille Université, CNRS, Marseille Cedex 20 F-13402, France
+present address: Laboratoire de Biodiversité et Biotechnologies Microbiennes (LBBM), Université de Perpignan Via Domitia, Sorbonne Université, CNRS, F-66860 Perpignan, France

#### S1 Enzyme preparation and biochemical characterization

##### Strains

*E. coli* strain DH5α (F-, endA1, hsdR17(rK-mK +), supE44, thi-1, λ-, recA1, gyrA96, relA1, Δ(argF-lacZYA)U169, φ80dlacZΔM15) was used as a host for the construction of recombinant plasmids. The cultures were routinely grown at 37 °C in Luria–Bertani (LB) medium containing 20 µg/mL gentamicin when needed. *Solidesulfovibrio fructosivorans* (formerly *Desulfovibrio fructosovorans)* (*Df*) strain MR400 (hyn::npt ΔhynABC) carrying a deletion in the [NiFe] hydrogenase operon[1] was grown anaerobically for 5 days at 37 °C in fructose/sulfate medium as previously described[2]. 50 µg/mL kanamycin was present routinely, and 20 µg/mL gentamicin was added only when cells harbored the expression vectors.

##### Plasmids construction for the production of Tc CODH 2 and variants in *Desulfovibrio fructosovorans*

The plasmid for the production of Tc CODH2 WT was constructed as described in [3]. Site directed mutagenesis was performed on the Tc CODH2 WT plasmid using primers carrying the desired nucleotide substitutions. The primers are shown in table S1. The entire plasmid was amplified by PCR with Pfu ultra HF from agilent (600380). Following amplification, the PCR products were treated with DpnI from NEB (R0176S) to digest the parental methylated plasmid DNA. The resulting nicked circular mutant plasmids were transformed into *E.coli DH5α* competent cells from NEB (C2987H). Positive clones were screened, verified by Sanger sequencing, and introduced into the *Df* strain MR 400 by electrotransformation. Exponentially growing cultures were harvested by centrifugation, washed several times with sterile cold water and resuspended at high density. Then mutant plasmids were introduced into the *Df* strain MR400 by electroporation using Gene pulser apparatus (Bio-Rad) with optimized parameters for this strain (with 1 mm electroporation cuvettes: 1.5 Kv, 25 µF, 400 ohms). Immediately after the electric pulse, cells were recovered in fructose sulfate medium, incubated under anaerobic conditions and after 5 hours the antibiotics were added to the cultures. After growth, 3 successive subcultures were subsequently carried out. The proteins were purified as described in [4] except that cells were disrupted in the glovebox by sonication or with Emulsiflex (Avestin Inc).

| Mutations | Forward primer (5'->3') | Reverse primer (5'-> 3') |
|---|---|---|
| I563E | AAAAGGCAGTCTCAGAAGGAACATACTT | AAGTATGTTCCTTCTGAGACTGCCTTTT |
| I563F | AAAAGGCAGTCTCATTTGGAACATACTT | AAGTATGTTCCAAATGAGACTGCCTTTT |
| I563Q | AAAAGGCAGTCTCACAAGGAACATACTT | AAGTATGTTCCTTGTGAGACTGCCTTTT |
| I563G | AAAAGGCAGTCTCAGGAGGAACATACTT | AAGTATGTTCCTCCTGAGACTGCCTTTT |
| I563A | AAAAGGCAGTCTCAGCAGGAACATACTT | AAGTATGTTCCTGCTGAGACTGCCTTTT |
| I563R | AAAAGGCAGTCTCACGAGGAACATACTT | AAGTATGTTCCTCGTGAGACTGCCTTTT |
| I563L | AAAAGGCAGTCTCACTGGGAACATACTT | AAGTATGTTCCCAGTGAGACTGCCTTTT |
| I563W | AAAAGGCAGTCTCATGGGGAACATACTT | AAGTATGTTCCCCATGAGACTGCCTTTT |
| F322H | ATAGCGGGAAGCCATCTGATGCAGGAACT | AGTTCCTGCATCAGATGGCTTCCCGCTAT |
| F322S | ATAGCGGGAAGCAGCCTGATGCAGGAACT | AGTTCCTGCATCAGGCTGCTTCCCGCTAT |

Table S1:List of primers used to generate mutations in this study.

#### Solution activity assays

The CO oxidation activity of *Tc* CODH 2 (WT and mutants) was measured in a glove box (Jacomex, filled with $N_2$, $O_2$ < 4 ppm) using a Varian Cary 50 spectrophotometer with a probe of 1 cm optical length. The CO oxidation activity was monitored at 37 °C by following the reduction of methyl viologen (MV) over time at 604 nm ($\varepsilon = 13.6\ mM^{-1}\ cm^{-1}$) in a plastic cuvette. Before recording the CO oxidation activity, the enzyme was diluted to a final concentration of 0.44 µM in 0.1 M Tris-HCl buffer, pH 8. To start the reaction, 5 µL of the enzyme solution were injected in a magnetic-stirred 1 mL cuvette containing 0.1 M Tris-HCl buffer (pH 8), 2.4 mM MV, 10 µM NaDT and 25 µM CO (25 µL of a CO-saturated solution injected just before the addition of the enzyme).

#### Elemental analysis

We determined the nickel and iron contents of the protein samples (500 µL of 5–10 µM protein solution) by ICP optical emission spectrometry (ICP-OES) using an iCAP 6000 spectrometer (ThermoFisher Scientific). Some of the variants were produced in too low quantities to be able to determine the metal contents.

#### Sequence alignment

The sequence alignment was generated based on the following accession codes:

Dv CODH: AAS96571.1
Rr CODH: AAC45123.1
Tc CODH1: EEB74637.1
Tc CODH2: EEB72960.1
To CODH: WP_012571978.1
Ch CODH1: P59934.3
Ch CODH2: ABB15588.1

Ch CODH3: 7ZKJ_A
Ch CODH4: 6ELQ_B
Mt ACS CODH: WP_053104382.1
Cae ACS CODH: 6YTT_B
Ci CODH: AHZ96929.1

### S2 Electrochemical experiments

Electroactive films of *Tc* CODH 2 (WT and mutants) were prepared by simply depositing 0.3 µL of enzyme stock solution (10–30 µM in 0.1 M Tris-HCl buffer, pH 8) onto a pyrolytic graphite edge rotating disc electrode (PGE-RDE, 2.5 mm diameter). All electrochemical experiments were performed in a standard three-electrode cell filled with 2 mL of the buffer solution of choice, placed inside a Jacomex glove box filled with $N_2$ ($O_2$ < 4 ppm). A platinum wire was used as the counter electrode, and a saturated calomel electrode (SCE, from Radiometer Analytical, France) as the reference, placed in a distinct compartment filled with 0.1 M NaCl. All electrode potentials are reported with respect to the standard hydrogen electrode (SHE). The electrodes were connected to an Autolab PGSTAT128N potentiostat (Metrohm, The Netherlands), controlled through the software GPES. The working electrode was rotated using an OrigaTrod electrode rotator (Origalys, France). All experiments were carried out by rotating the working electrode at 4000 rpm to minimize depletion of the substrate at the electrode surface. The temperature of the water-jacket surrounding the electrochemical cell was kept constant during the experiments at 25 °C.

All electrochemical experiments to determine the catalytic bias, the $K_M$ for CO and $CO_2$, the product inhibition and $O_2$ inhibition for *Tc* CODH 2 (WT and mutants) were performed like described in ref [3]for the wild type of *Tc* CODHs. However, for the experiments for the determination of the $K_M$ for $CO_2$ and the product inhibition of $CO_2$ by CO, we realized that our previous data analysis was flawed, because it did not take into consideration the inhibition by the CO produced by the reaction but not evacuated yet. Figure S1 shows the data, and we now describe their interpretation.

We assume that the concentration of $CO_2$ is not impacted by mass transport (this is justified considering that the concentrations of $CO_2$ involved are several orders of magnitude greater than those of CO), and that the concentration of CO at the electrode is given by the classical hydrodynamic electrode equation:

$$[\mathrm{CO}]^0 = [\mathrm{CO}]^\infty - \frac{J_{\mathrm{CO}}}{m}$$

in which $[\mathrm{CO}]^\infty$ is the concentration of CO far away from the electrode, $[\mathrm{CO}]^0$ the concentration of CO at the electrode, $J_{\mathrm{CO}}$ the flux of CO towards the electrode, and *m* the mass-transport coefficient of CO. The flux of CO towards the electrode is the opposite of the flux of $CO_2$ towards the electrode, given by:

$$J_{\mathrm{CO}_2} = -J_{\mathrm{CO}} = \Gamma \frac{k_{cat}[\mathrm{CO}_2]}{K_M \left(1 + \frac{[\mathrm{CO}_2]}{K_M} + \frac{[\mathrm{CO}]^0}{K_i}\right)}$$

in which $\Gamma$ is the electroactive coverage, $k_{cat}$ the maximum turnover frequency for the $CO_2$ reduction, $K_M$ the Michaelis constant for $CO_2$, $K_i$ the product inhibition constant of $CO_2$ by CO. This can be rearranged thus:

$$J_{\mathrm{CO}_2} = \frac{J^{eff}}{1 + \frac{[\mathrm{CO}]^0}{K_i^{eff}}}$$

with the following definitions:

$$K_i^{eff} = K_i \left(1 + \frac{[\mathrm{CO}_2]}{K_M}\right)$$

$$J^{eff} = \Gamma \frac{k_{cat}[\mathrm{CO}_2]}{K_M + [\mathrm{CO}_2]}$$

Solving for the concentration of CO at the electrode gives the following concentration:

$$[\mathrm{CO}]^0 = \frac{1}{2}\left([\mathrm{CO}]^\infty - K_i^{eff} + \sqrt{\left(K_i^{eff} + [\mathrm{CO}]^\infty\right)^2 + 4\frac{J^{eff}K_i^{eff}}{m}}\right)$$

The flux can be expressed as a function of the maximum current $i_{max}$ and *F*, the Faraday constant, *n* (=2), the number of electrons involved in the catalytic reaction and *A* the electrode area, or alternatively using μ, the product of the surface coverage by the catalytic efficiency:

$$i_{max} = nFA\Gamma k_{cat} = nFAK_M\Gamma\frac{k_{cat}}{K_M} = nFAK_M\mu$$

To compute the current, we used the following ruby function that takes as arguments `s` the concentration of $CO_2$, `p_inf` the concentration of CO infinitely away from the electrode (i.e. the concentration of injected CO), `km` the $K_M$ for $CO_2$, `ki` the product inhibition constant $K_i$, `imax` the maximum current $i_{max}$ and `nFAm` the product $nFAm$.

```
def inhib_transport(s, p_inf, km, ki, imax, nFAm)

  i_eff = imax * s/(s + km)

  ki_eff = ki * (1 + s/km)

  p_real = 0.5*(p_inf - ki_eff + ((ki_eff + p_inf)**2 + 4 * i_eff *
ki_eff/nFAm)**0.5)

  return i_eff/(1 + p_real/ki_eff)

end
```

This function was used with the following command to fit the data in QSoas:

```
mfit-arb inhib_transport(co2,co,km,ki,nFAmu*km,nFAm)+io
/with=co2:1,co2;co:1,exp 0,1,2
```

This command directly uses $nFA\mu$ as a fit parameter, because it is better defined than the limiting current in conditions where the value of $K_M$ is very large.

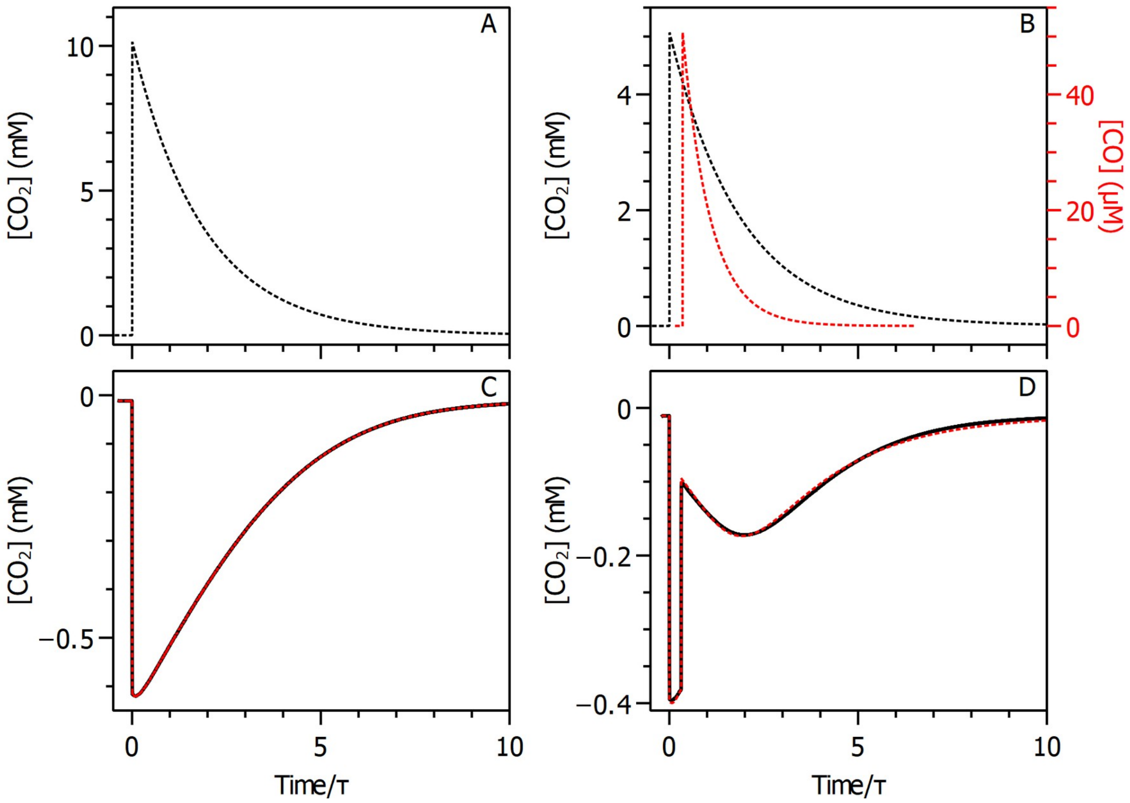


Fig. S1. Chronoamperometric measurements of the Michaelis and product inhibition constants relative to $CO_2$. The time is normalized by the characteristic time of departure of $CO_2$ from solution. A) Concentration of $CO_2$ in the cell vs. time. B) Black line: concentration of $CO_2$ in the cell vs. time vs. the left axis. Red line: concentration of CO in the cell vs. time vs. the right axis. C) Black line: current observed with films of Tc CODH2 F322S subjected to a single injection of 10 mM $CO_2$ as in panel A). D) Black line: current observed with films of Tc CODH2 F322S subjected to an injection of 10 mM $CO_2$, followed by an injection of 47 µM CO as in panel B). The $CO_2$ departure time constants are 58 s and 64 s in C and D respectively, the CO departure time constant in D is 40 s. Experimental conditions: T = 25 °C, pH 7, electrode rotation rate ω = 4000 rpm, E = −0.66 V vs. SHE. Red dashed lines in C) and D): fit of the chronoamperograms using Eq. X, C) and D) were fitted together using the `mfit-arb` command in QSoas. Parameters of the fits: C) $K_i$ = 10 µM, $K_M$ = 2825 µM nFAµ = -0.292 nA/µM, nFAm = 0.15 µA/µM, i0 = −10.66 nA, [CO]0 = 0 µM, τ CO = 40 s, pH = 6.37, pKa = 6.4, τ CO2 equilibrium = 5 s, [CO2]0 = 4501 µM, [HCO3]0 = 5400 µM, τ CO2 = 58. D) Ki = 10 µM, Km = 2825 µM nFAµ = -0.247 nA/µM, nFAm = 0.15 µA/µM, i0 = −10.75 nA, [CO]0 = 47 µM, τ CO = 40 s, pH = 6.44, pKa = 6.4, τ CO2 equilibrium = 5 s, [CO2]0 = 2076 µM, [HCO3]0 = 2850 µM, τ CO2 = 64.

### S3 Definition of the $O_2$-sensitivity parameters

The apparent inhibition constants: $K_i^{imm}$, $K_i^{dep}$, and $K_i^{red}$ from each series of experiments are determined from the values imm, dep and red, which are defined as the remaining active fraction shortly after $O_2$ injection (imm), after $O_2$ departure (dep), and after exposure to low potential (red) according to the following equations:

$$imm = \frac{i_{3,50}/i_3}{\left(i_{1,50}/i_1 + i_{2,50}/i_2\right)/2}$$

$$dep = \frac{i_4 / i_3}{\left(i_2 / i_1 + i_6 / i_5\right) / 2}$$

$$red = \frac{i_5 / i_3}{i_3 / i_1}$$

In which $i_1$ through $i_6$ are the currents 2 s after the 1st through 6th CO injections and $i_{1,50}$ through $i_{3,50}$ are the currents 50 seconds after the respective CO injection as shown in main text figure 4. In each equation the numerator is used to calculate the remaining active fraction at the point of interest while the denominator is used to correct for film loss during the experiment. Each of the three active fractions was determined for a range of $O_2$ concentrations as in main text Figure 5, so that the point at which 50% of the activity remains can be determined for each, giving $K_i^{imm}$, $K_i^{dep}$, and $K_i^{red}$.

### S4 Alphafold predictions

The structures of *Tc* CODH 2 WT (as control) and all the I563 mutants were predicted using alphafold 3. In particular, 5 independent runs of alphafold were performed, and only the best ranked structure per each run was used. The metallic clusters D, B and C of *Tc* CODH 2 WT (PDB 6T7J) were inserted in the alphafold models using pymol.

### S5 CAVER prediction of the channels

The hydrophobic channels were computed using CAVERanalyst64 with the following configuration:

probe_radius: 0.9
shell_radius: 3
shell_depth: 6

First, we computed the tunnels from the crystal structure of *Tc* CODH 2 WT (PDB 6T7J), using the exo Fe of the C cluster as starting point. Then we computed the channels for the 5 alphafold models of *Tc* CODH 2 WT and all the I563 mutants. To avoid possible artifacts, per every set of 5 alphafold models, we only kept the models whose predicted channels matched the ones computed with *Tc* CODH 2 WT crystal structure. After this screening we have:

- Tc2 PDB 6t7j: 13 tunnels
- Tc2 WT: 5 models (16 to 20 tunnels)
- Tc2 I563L: 5 models (13 to 19 tunnels)
- Tc2 I563Q: 4 models (13 to 19 tunnels)
- Tc2 I563A: 4 models (16 to 22 tunnels)
- Tc2 I563E: 5 models (15 to 18 tunnels)
- Tc2 I563F: 5 models (14 to 19 tunnels)
- Tc2 I563W: 4 models, all having only 1 of the 2 WT channels (7 to 13 tunnels)

For all the above models, CAVER prediction counted many channels (between 13 and 22 depending on the mutant and on the model), differing in the final parts, but being all very similar in the part closer to the active site.
Because our interest is focused on the branching point, where the one channel leaving the active site splits into two branches (close to the 563 position), we selected the 2 best ranked channels representative of the 2 branching per every selected model.

Statistics that generated figure 6 in the main text:

We computed the distance of each point of the channels from the starting point (exo Fe) with the following formula: `sqrt((x-x0)**2+(y-y0)**2+(z-z0)**2)`. In figure 7 is plotted the average radius as a function of distance computed from the 2 best channels of each selected model of all the mutants.

The width profile of the two channels is shown in figure S2 as a function of the distance from the position of the unique iron, $Fe_u$, for all of the I563 variants described in this article. Only the first 15 Å are shown, which corresponds to the common part and the beginning of each branch. On each panel, the gray dashed lines correspond to the channel observed in the crystal structure of the WT, while the black solid lines are determined from the AlphaFold models. The top panel shows that the channel features CAVER predicts from the AlphaFold model are very similar to the crystal structure of the WT. Unsurprisingly, the channels of the I563L variant barely differ from those of the WT. The I563A, I563F and I563W variants all present a significant broadening of the common region of the channels with, for the F and W substitutions, a significant narrowing further away from the C cluster. On the other hand, the channels of the I563E and I563Q variants are constricted in the shared region, also with some narrowing further in the channels.

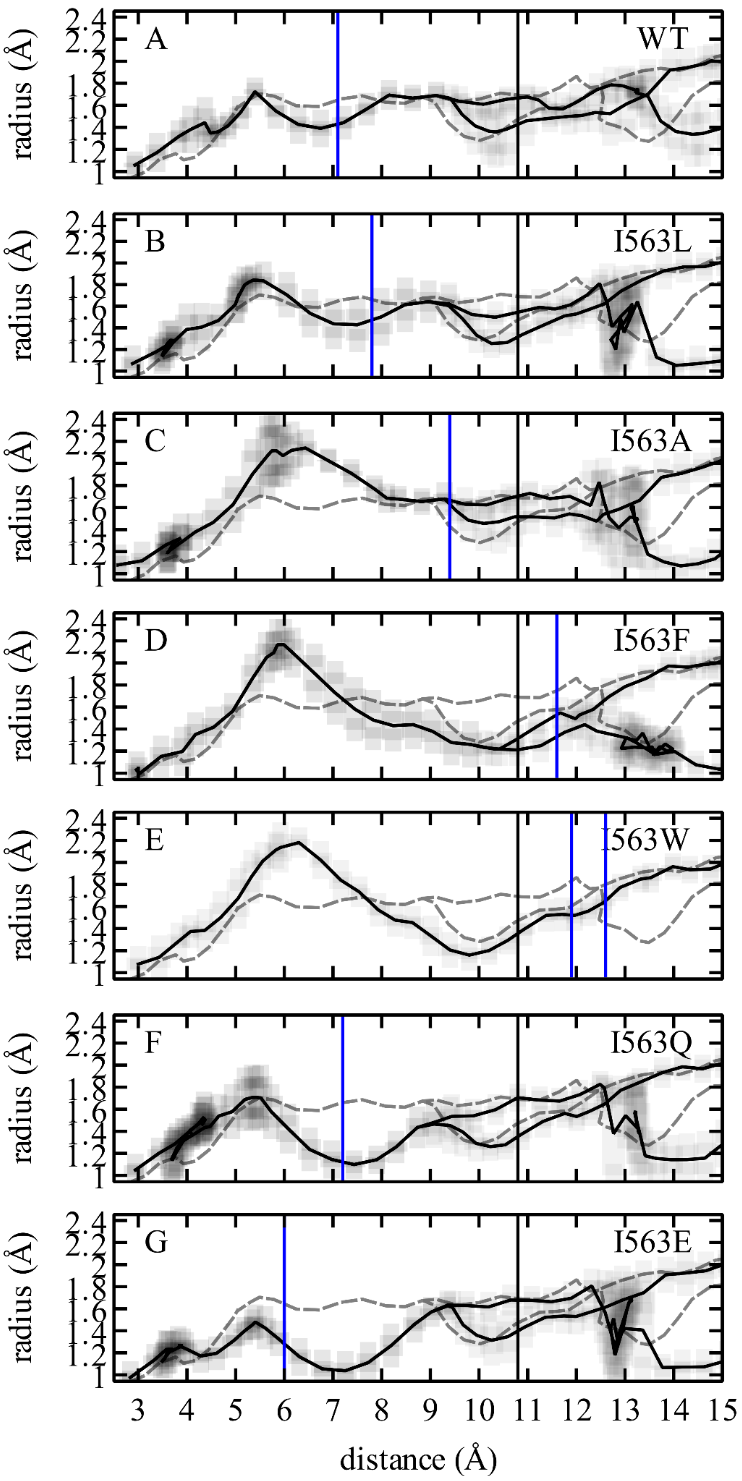


Figure S2. Radius of the 2 tunnels as a function of the distance from $Fe_u$. In every panel, the gray dashed lines are the radius of the tunnels computed from the crystal structure of *Tc* CODH2 WT (PDB 6T7J) and the solid black lines are the average radius of the tunnels computed from different AlphaFold models. The number of AlphaFold models used to compute the channel radius is 4 (panels C, F and E) or 5 (panels A, B, D and G), see the SI sections S4 and S5. The standard deviation is plotted as a shaded gray area. Vertical black lines mark the distance between $Fe_u$ and the C alpha of the amino acid at position 563. Vertical blue lines mark the distance between $Fe_u$ and the “last” carbon of the side chains of the amino acid at position 563. The I563W variant has 2 blue lines since, out of the 5 AlphaFold models, 2 slightly different positions of tryptophan were modeled.

### S6 Further correlation plots

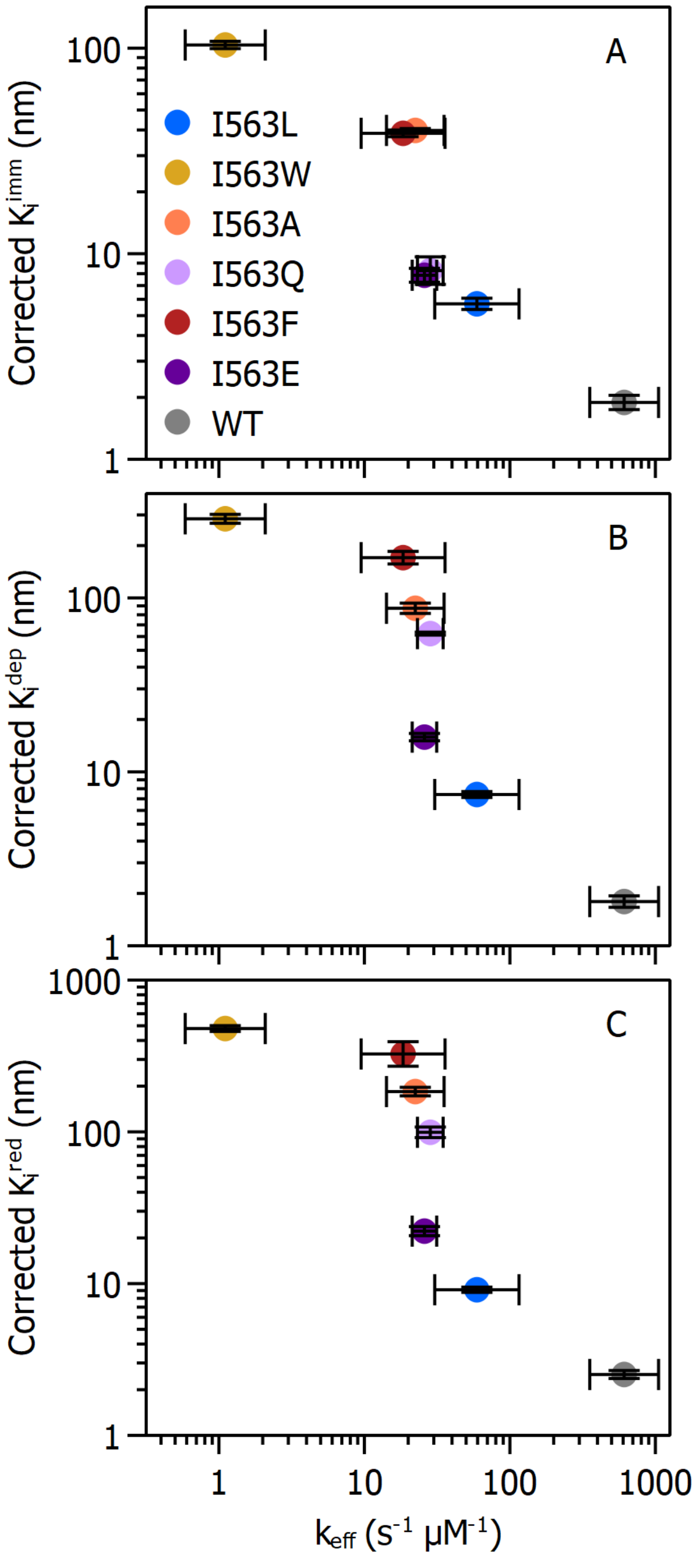


Figure S3. Plot of the corrected values Kiimm, Kidep and Kired as a function of the catalytic efficiency, for all the variants. The apparent inhibition constants are corrected as expected for a competitive inhibition, i.e. $K_i^{corr} = K_i/(1 + [CO]/K_M)$. The trend is the inverse of that of $k_{O2}$: the lower the $k_{O2}$, the more the enzyme is able to resist a high quantity of $O_2$, thus the higher the corresponding apparent inhibition constant.

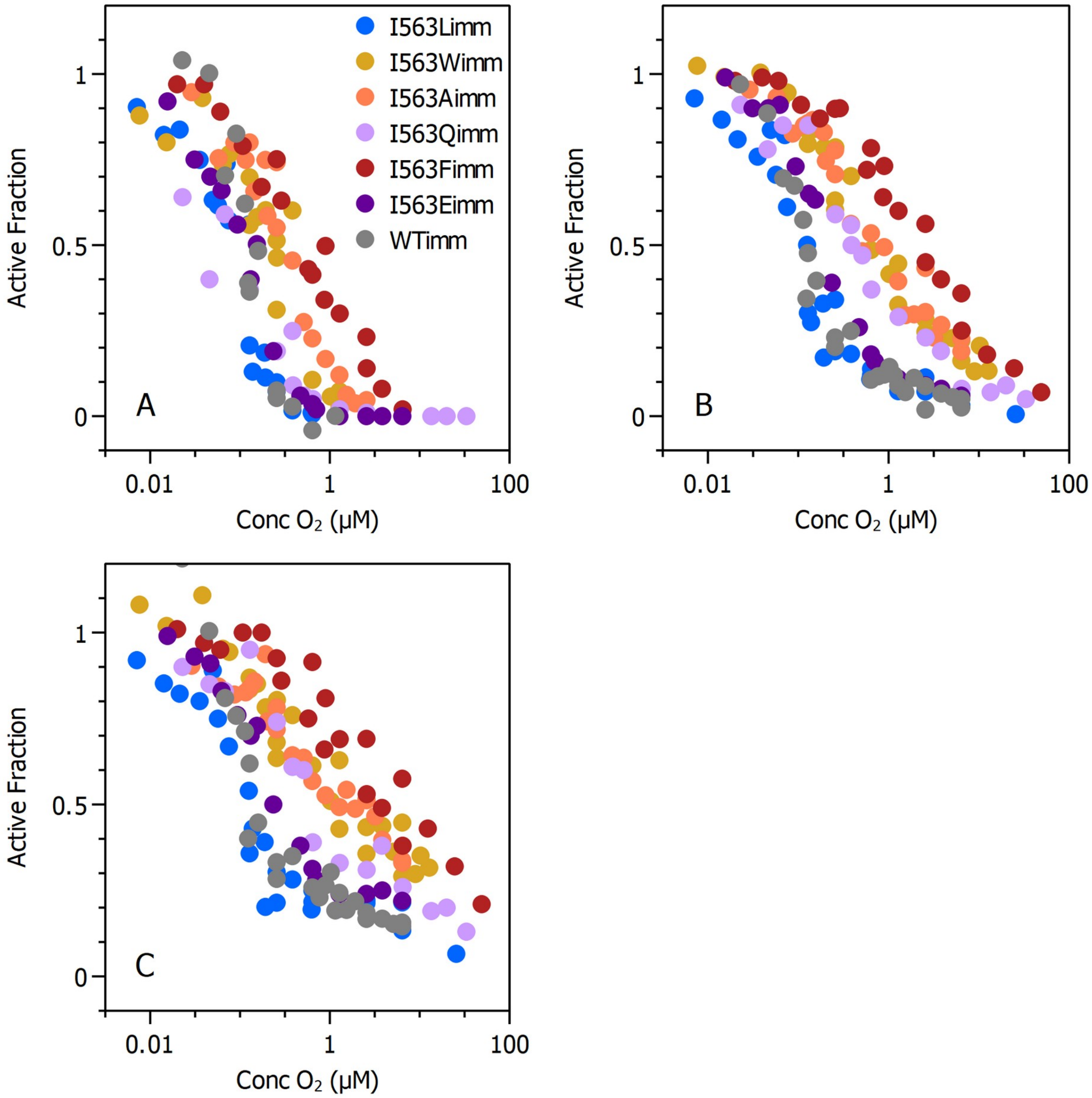


Figure S4. Plot of the remaining active fraction immediately after exposure to oxygen (A), after oxygen departure (B), and after a reductive poise © as deduced from the 7-injections experiment in PFV as plotted in Figure 6. Here the imm, dep, and red values for the WT and all variants are plotted together to permit easier comparison.
